# The PfAP2-HS transcription factor protects malaria parasites from febrile temperatures

**DOI:** 10.1101/2021.03.15.435375

**Authors:** Elisabet Tintó-Font, Lucas Michel-Todó, Timothy J. Russell, Núria Casas-Vila, David J. Conway, Zbynek Bozdech, Manuel Llinás, Alfred Cortés

## Abstract

Periodic fever is the most characteristic clinical feature of human malaria^1-3^, but how parasites survive febrile episodes is not known. While *Plasmodium* spp. genomes encode a full complement of chaperones^4^, they lack an ortholog of the conserved transcription factor HSF1, which in most eukaryotes activates the expression of key chaperones upon heat shock (HS)^5-8^. Here we identified PfAP2-HS, a transcription factor of the ApiAP2 family^9-11^, as the key regulator of the *P. falciparum* protective HS response. The PfAP2-HS-dependent HS response is largely restricted to rapid activation of *hsp70-1*, the predominant direct target of PfAP2-HS, and *hsp90*. Deletion of PfAP2-HS dramatically reduced HS survival and also resulted in severe growth defects at 37°C, but not at 35°C, and increased sensitivity to imbalances in protein homeostasis (proteostasis) produced by artemisinin, the current frontline antimalarial drug^12,13^. These results demonstrate that PfAP2-HS contributes to general maintenance of proteostasis and drives a rapid chaperone-based protective response against febrile temperatures. While several ApiAP2 transcription factors regulate life cycle transitions in malaria parasites^11,14,15^, PfAP2-HS is the first identified *Plasmodium* transcription factor that controls a protective response to a within-host environmental challenge.

---

A temperature increase of just a few degrees above the optimal growth temperature can cause aberrant protein folding and aggregation, resulting in a proteostasis imbalance that can lead to cell-cycle arrest or even cell death^16^. To counteract the effect of high temperatures and other proteotoxic conditions, living organisms use a complex array of molecular chaperones that assist protein refolding and prevent non-specific aggregation^16,17^. This is especially important for human malaria parasites, which are exposed to periodic fever episodes during blood stages. In *P. falciparum* infections, which are responsible for the most severe forms of human malaria, fever typically occurs on alternate days (tertian fever). This reflects the ∼48 h duration of the asexual intraerythrocytic development cycle (IDC), during which parasites progress through the ring, trophozoite and multinucleated schizont stages. Schizont rupture and release of new invasive merozoites trigger fever episodes, an important component of the innate immune response that limits the total parasite burden^2,3,18^.

To understand the molecular basis of HS resistance in *P. falciparum*, we first analysed parasite lines that we had previously selected with periodic HS over five consecutive rounds of the IDC^19^ (Supplementary Fig. 1a). Although the parental parasite line (3D7-A) appeared to have lost the ability to withstand febrile temperatures during growth in culture, its ability to quickly re-adapt to HS pressure suggested that this line contained a selectable subpopulation of parasites resistant to HS, which could have either a genetic or an epigenetic basis. Transcriptomic analysis across the full IDC under basal conditions (no HS) of two independently selected lines (3D7-A-HS r1 and r2) and non-selected cultures maintained in parallel (3D7-A r1 and r2) failed to identify any basal transcript level differences that could explain the HS resistance phenotypes (Supplementary Fig. 1b-d). Therefore, we sequenced the genomes of these lines, which revealed a novel single nucleotide polymorphism (SNP) that was predominant in non-selected cultures, but virtually absent after HS selection (Fig. 1a-b, Supplementary Table 1). The mutation is also absent from two other 3D7 stocks in our laboratory and from the 3D7 reference genome, indicating that it arose spontaneously in the 3D7-A stock during culture. This SNP results in a premature STOP codon (Q3417X) in the gene PF3D7_1342900, which encodes a putative transcription factor of the ApiAP2 family^9-11^ that we termed PfAP2-HS. PfAP2-HS has three AP2 domains (D1-D3), and the Q3417X mutation results in a truncated protein that lacks D3 (PfAP2-HSΔD3) (Fig. 1c). Therefore, adaptation to HS involved elimination of parasites that expressed truncated PfAP2-HS, consistent with a role for this protein in the HS response. In support of this idea, the first AP2 domain (D1) of PfAP2-HS was previously reported to recognize *in vitro* a DNA motif termed G-box^10^, which is enriched in the upstream region of some HS protein (HSP) chaperone genes^20^.

**Fig 1.**
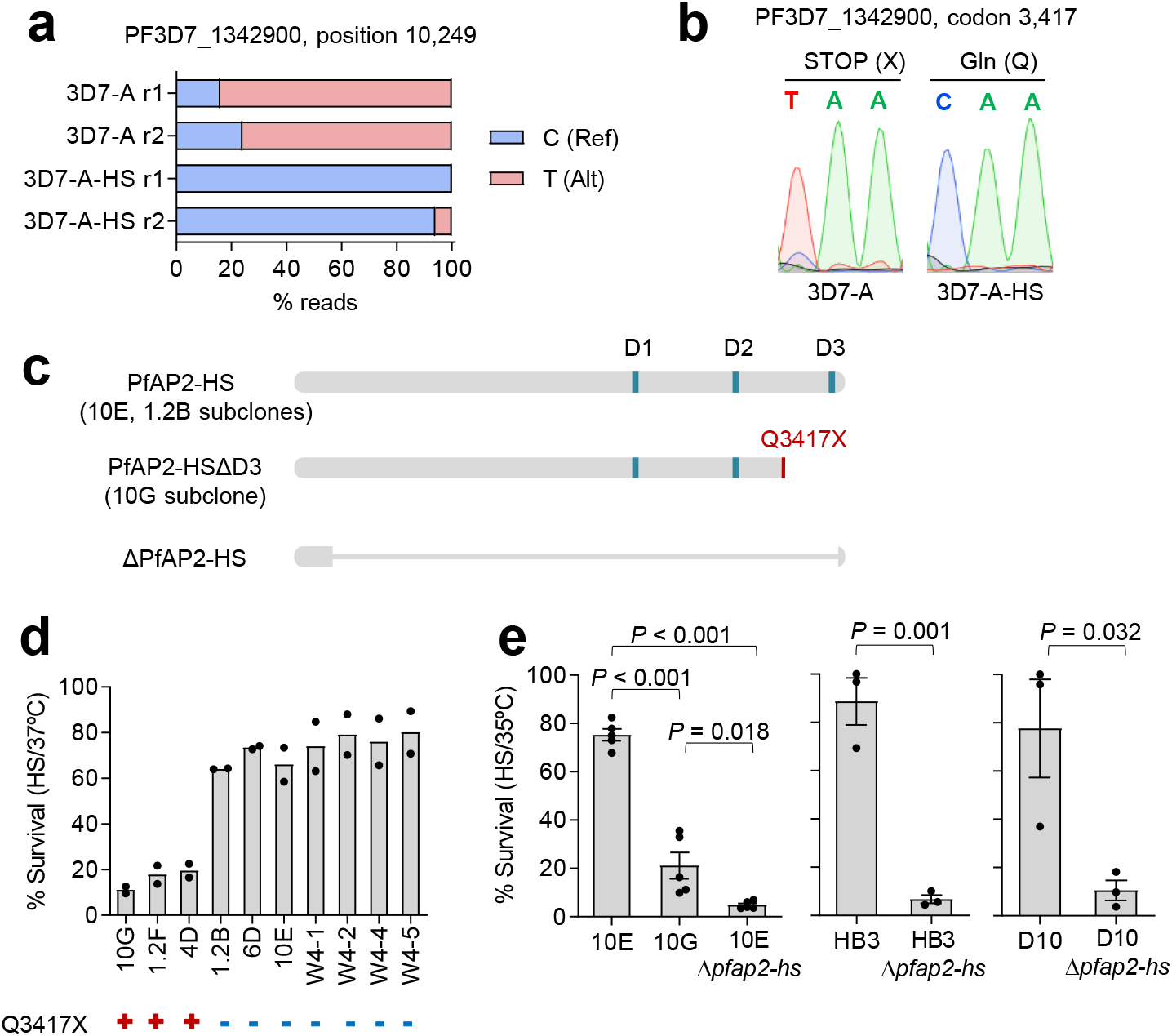
A premature STOP codon in PfAP2-HS determines HS sensitivity. **a**, Proportion of Illumina reads with or without a nonsense mutation in *pfap2-hs* in two independent HS-adapted cultures (3D7-A-HS r1 and r2) and their controls (3D7-A r1 and r2). **b**, Sanger sequencing confirmation of the mutation. **c**, Schematic of wild-type PfAP2-HS, PfAP2-HS_ΔD3 and ΔPfAP2-HS. The position of the AP2 domains is indicated (D1-3). **d**, HS survival at the trophozoite stage of 3D7-A subclones carrying or not the Q3417X mutation (mean of *n*=2). **e**, HS survival of tightly synchronized wild-type, 10G and Δ*pfap2-hs* cultures. Values are the mean and s.e.m. of *n*=5 (10E and 10G lines) or *n*=3 (HB3 and D10 lines).

To test the involvement of PfAP2-HS in HS resistance, we used a HS survival assay with a 3 h HS at 41.5°C^19^ at the mature trophozoite stage, because maximal survival differences between HS sensitive and resistant parasite lines were observed when exposing parasites at this stage (Supplementary Fig. 1e). The analysis of a collection of 3D7-A subclones revealed that all subclones with the Q3417X mutation show a HS-sensitive phenotype, whereas subclones with the wild-type allele show a resistant phenotype (Fig. 1d).

To further characterize PfAP2-HS, we disrupted the entire gene using CRISPR-Cas9 technology. After several unsuccessful attempts with different 3D7 subclones at 37°C (the physiological temperature for *P. falciparum*), we reasoned that PfAP2-HS may play a role in regulating the expression of chaperones under basal conditions, in addition to being necessary for HS survival. Therefore, we attempted to knock out the gene in cultures maintained at 35°C, as mild hypothermia is expected to reduce protein unfolding and favour proteome integrity^21,22^. Indeed, at 35°C, knockout of *pfap2-hs* was readily achieved in both the HS-resistant 10E and the HS-sensitive 10G subclones of 3D7-A (10E_Δ*pfap2-hs* and 10G_Δ*pfap2-hs* lines) (Fig. 1c, Supplementary Fig. 2a). Deletion of *pfap2-hs* resulted in severely increased sensitivity to HS, with a level of HS survival below that of parasites expressing PfAP2-HSΔD3. Deletion of the gene in two parasite lines of unrelated genetic background, HB3 and D10, also resulted in a major reduction in HS survival (Fig. 1e).

To define the PfAP2-HS-dependent and independent HS response, we conducted a transcriptomic time-course analysis of the 10E (wild-type PfAP2-HS), 10G (PfAP2-HSΔD3) and 10E_Δ*pfap2-hs* lines during and after HS (Fig. 2a, Supplementary Fig. 3a, Supplementary Table 2). Hierarchical clustering based on changes relative to cultures not exposed to HS revealed only three genes in the full genome with a rapid increase in transcript levels upon HS that occurred in 10E but not in 10G or 10E_Δ*pfap2-hs* (cluster I). This cluster comprises a gene of unknown function (PF3D7_1421800), the cytoplasmic *hsp70* (*hsp70-1*; PF3D7_0818900) and *hsp90* (PF3D7_0708400) (Fig. 2a). These two chaperone-encoding genes contain in their regulatory regions the best two matches in the full genome for a tandem G-box^10,20^ (Supplementary Fig. 3b), suggesting an important role for this tandem arrangement. The strongest transcriptional response was observed for *hsp70-1* (∼16-fold increase versus ∼4-fold for *hsp90*). To validate the observation that rapid activation of cluster I genes upon HS depends on PfAP2-HS and requires its D3, we used *pfap2-hs* knockout parasite lines of different genetic backgrounds and several 3D7-A mutant subclones expressing PfAP2-HSΔD3 (Supplementary Fig. 4). In all knockout and mutant lines examined, the *hsp70-1* response to HS was delayed and of much lower magnitude. These experiments also confirmed that the *hsp90* response is weaker than the *hsp70-1* response, and is delayed in PfAP2-HS mutants.

**Fig 2.**
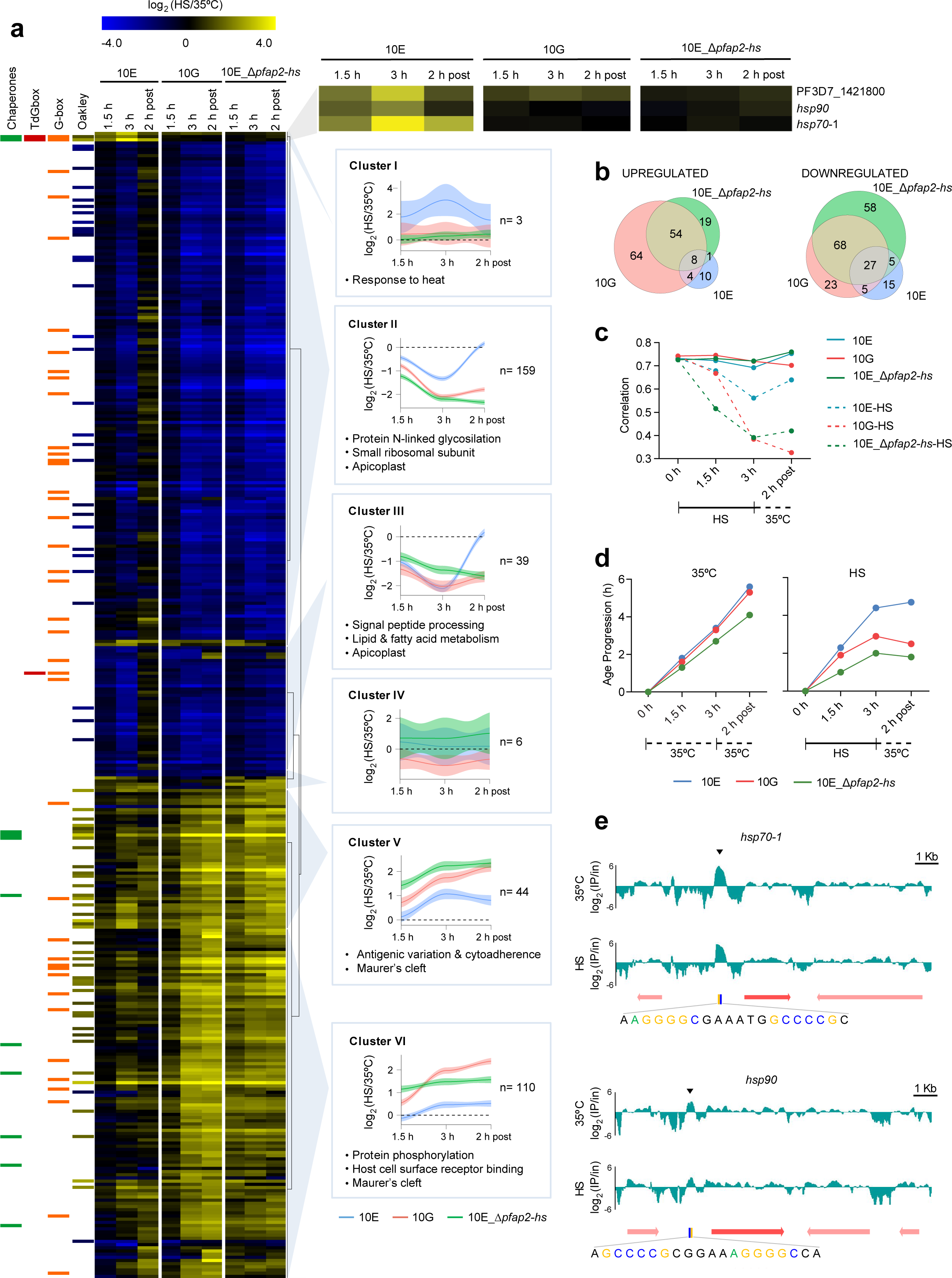
Global transcriptional alterations upon HS. **a**, Hierarchical clustering of genes with altered transcript levels (≥4 fold-change at any of the time points analysed) during (1.5 and 3 h) or after (2 h post) HS. Values are the log_2_ of the expression fold-change in HS versus control cultures. 13 genes had values out of the range displayed (actual range: -4.78 to +4.93). For each cluster, mean values (with 95% confidence interval) for the genes in the cluster and representative enriched GO terms are shown. Columns at the left indicate annotation as chaperone^4^, presence of the G-box^10^ or tandem G-box (TdGbox) in the upstream region, and log_2_ fold-change during HS in a previous study^24^ (Oakley). **b**, Venn diagrams of the genes altered upon HS in the three parasite lines. **c**, Pearson correlation of the genome-wide transcript levels of each culture versus the most similar time point of a high-density time-course reference transcriptome^23^. **d**, Age progression during the assay, statistically estimated^53^ from the transcriptomic data. **e**, ChIP-seq analysis (representative of *n*=5 for 35°C and of *n*=3 for HS) of HA-tagged PfAP2-HS. The log_2_-transformed ChIP/input ratio at the *hsp70-1* and *hsp90* loci is shown. The position of the tandem G-box is indicated.

Genes in other clusters (II-VI) of the transcriptomic analysis showed changes in expression during HS that did not depend on PfAP2-HS, as they occurred even in the knockout line (Fig. 2a). Indeed, a larger number of genes with altered transcript levels was identified in 10G and 10E_Δ*pfap2-hs* than in 10E (Fig. 2b). Furthermore, the alterations persisted 2 h after HS in both HS-sensitive lines, whereas in 10E the majority of genes returned to basal expression levels (Fig. 2a). This indicates that many of these alterations reflect unresolved cell damage or death, and in 10E the rapid PfAP2-HS-dependent response limited cell damage and the subsequent changes in the expression of clusters II-VI genes. Indeed, after HS the transcriptome of the 10G and 10E_Δ*pfap2-hs* lines showed a more pronounced deviation from a reference transcriptome^23^ than 10E (Fig. 2c). Global transcriptional analysis also revealed that HS resulted in delayed IDC progression, again more pronounced in 10G and 10E_Δ*pfap2-hs* (Fig. 2d).

In addition to genes reflecting cell damage, clusters II-VI likely include some genes that participate in the PfAP2-HS-independent HS response. In particular, clusters V-VI include several chaperone-encoding genes upregulated during HS, although at a later time point than cluster I genes (Fig. 2a). However, the expression of the majority of known *P. falciparum* chaperones^4^ was not altered by HS and, except for cluster I genes, the alterations occurred mainly in the mutant lines (Supplementary Fig. 5). To provide a clearer view of the wild-type HS response, we analysed changes upon HS in the 10E line alone. Overall, there was generally good concordance with the genes and processes altered upon HS described in a previous study using non-synchronized cultures^24^ (Supplementary Fig. 6, Supplementary Table 3). Altogether, we conclude that a number of genes are up or downregulated during HS and some may contribute to HS protection through PfAP2-HS-independent responses, but in the absence of the rapid PfAP2-HS-dependent activation of cluster I genes these responses are not sufficient to ensure parasite survival under febrile temperatures.

To determine the genome-wide occupancy of PfAP2-HS, we analysed a parasite line expressing endogenous HA-tagged PfAP2-HS (Supplementary Fig. 2d) using chromatin immunoprecipitation followed by sequencing (ChIP-seq). The predominant binding site of PfAP2-HS coincides with the position of the tandem G-box in the upstream region of *hsp70-1* (Fig. 2e, Supplementary Fig. 7, Supplementary Table 4). This is the only binding site with a median fold-enrichment >10 (ChIP versus input) that was consistently detected across five independent replicates, revealing an extremely restricted distribution of PfAP2-HS binding. Similar enrichment was observed between control and HS conditions by both ChIP-seq and ChIP-qPCR (Supplementary Fig. 7a,c), indicating that PfAP2-HS binds to this site constitutively and is activated *in situ* upon HS. This is similar to human HSF1, which in the absence of HS is bound to its target promoters in a paused state to ensure a rapid response^6^. Enrichment at the *hsp90* promoter also coincided with the position of the G-box but was much weaker and, in several replicates, a significant peak was not called. No enrichment was observed at the promoter of cluster I gene PF3D7_1421800 that lacks a G-box motif. The only other site consistently enriched for PfAP2-HS binding, albeit at much lower levels than *hsp70-1*, was the upstream region of the small nucleolar RNA *snoR04* (PF3D7_0510900) (Supplementary Fig. 7), which was not upregulated during HS.

Both Δ*pfap2-hs* lines of 3D7 origin (10E_Δ*pfap2-hs* and 10G_Δ*pfap2-hs*) showed severe temperature-dependent growth defects in the absence of HS. They grew at similar rates to the parental lines at 35°C, but their growth was markedly reduced at 37°C or 37.5°C (Fig. 3a). The D10 Δ*pfap2-hs* line also had clearly reduced growth at 37°C compared to 35°C, whereas the HB3 Δ*pfap2-hs* line did not (Supplementary Fig. 8a). Both 3D7 knockout lines also showed a reduced number of merozoites per schizont, especially at 37°C or 37.5°C (Fig. 3b), which partly explains the reduced growth rate. Additionally, even at 35°C, the duration of the IDC was ∼4 h longer in both knockout lines (Fig. 3c), which is reminiscent of the slower life cycle progression observed in parasites under proteotoxic stress^25^. In contrast, no growth rate or life cycle duration differences were observed between the 10G (PfAP2-HSΔD3) and 10E lines, indicating that D3 is necessary for HS survival but not for growth under nonstress conditions (Fig. 3a-c). Normal growth at 37°C but low HS survival was also observed in transgenic lines in which bulky C-terminal tags were added to the C-terminus of endogenous PfAP2-HS, suggesting interference of the tag with the function of D3, located only 18 amino acids from the end of the protein (Supplementary Fig. 2).

**Fig 3.**
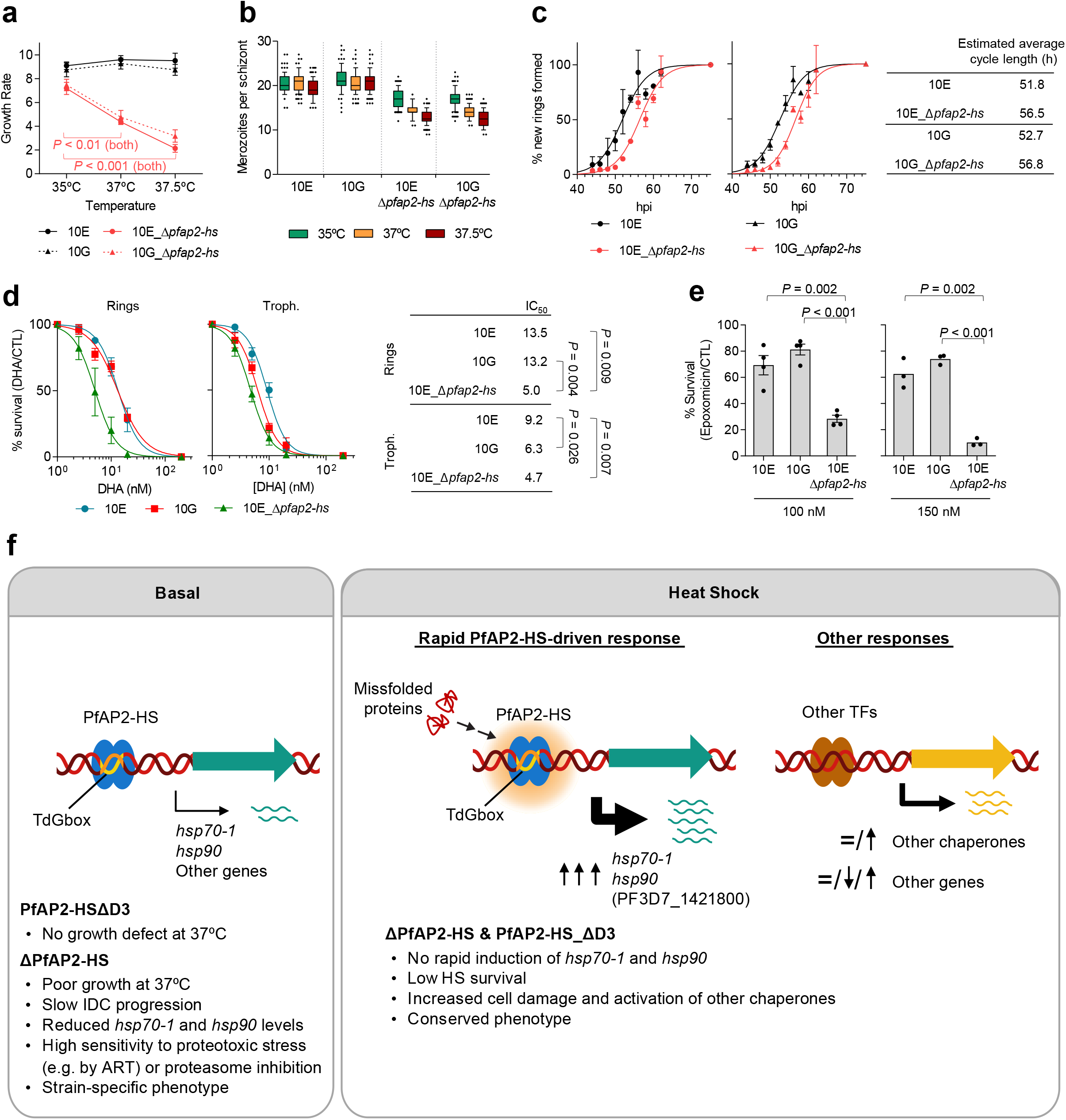
Phenotypic characterization of parasite lines lacking PfAP2-HS at non-febrile temperatures. **a**, Growth rate of Δ*pfap2-hs* and parental lines at different temperatures (mean and s.e.m. of *n*=4). **b**, Number of merozoites per schizont (median and quartiles box with 10-90 percentile whiskers). Values were obtained from 100 schizonts for each parasite line and condition. **c**, Duration of the asexual blood cycle. The cumulative percent of new rings formed at each time point is shown (mean and s.e.m. of *n*=2). **d**, Survival (%) after a 3 h dihydroartemisinin (DHA) pulse at the ring or trophozoite stage (mean and s.e.m. of *n*=3). **e**, Survival (%) after a 3 h epoxomicin pulse at the trophozoite stage. Values are the mean and s.e.m. of *n*=4 (100 nM) or *n*=3 (150 nM). **f**, Schematic model of the *P. falciparum* HS response, involving rapid upregulation of the expression of a very restricted set of key chaperones by PfAP2-HS. The PF3D7_1421800 gene (in brackets) shows PfAP2-HS-dependent increased transcript levels upon HS, but PfAP2-HS binding was not detected in its promoter, and it lacks a G-box. The main defects associated with PfAP2-HS deletion or truncation, under HS or basal conditions, are listed.

Genome-wide sequence analysis has recently revealed that non-sense mutations commonly emerge in *pfap2-hs* during the adaptation to culture conditions^26,27^, whereas they are virtually absent from clinical isolates (in the www.malariagen.net/data/pf3k-5 dataset^28^, only one out of >2,500 isolates carries a high-confidence SNP resulting in a premature stop codon). This indicates a likely essential role for PfAP2-HS in human infections, and relaxed selection against loss-of-function when parasites are not exposed to febrile conditions. We chose for further analysis a culture-adapted isolate carrying a mutation that results in truncation of PfAP2-HS before its first AP2 domain^26^ (monoclonal Gambian Line 1, PfAP2-HS_ΔD1-3), and found that a single HS resulted in strong selection against parasites carrying the mutation. In contrast, there was relatively weak selection against mutants during culture either at 35°C or 37°C, as the prevalence of the mutation only decreased from ∼50% to ∼20% after culturing for 23 generations (Supplementary Fig. 8b-d). Consistent with these results, a subclone carrying the mutation (1H) was more sensitive to HS than a wild-type subclone (4E), but both showed no measurable difference between growth at 35°C or 37°C (Supplementary Fig. 8e-g). Together, these results indicate that PfAP2-HS is essential for HS survival in all the genetic backgrounds tested. However, PfAP2-HS is necessary for normal progression through the IDC at 37°C only in some genetic backgrounds (i.e., 3D7 and D10), whereas in others (HB3 and the Gambian isolate) it is not essential.

To gain insight on the molecular basis of the growth defects of some of the knockout lines, we compared the trophozoite transcriptome of 10E_Δ*pfap2-hs* with that of the parental 10E under basal (no HS) conditions. This revealed only a small set of genes with a ≥2 fold-decrease in transcript levels, which included *hsp70-1*, the direct PfAP2-HS target *snoR04* RNA and several genes mainly involved in ribosome formation. Transcript levels for *hsp90* were also reduced (<2 fold-decrease) in the knockout line (Supplementary Fig. 9a-b). Lower transcript levels of *hsp70-1* and *hsp90* under basal conditions in 10E_Δ*pfap2-hs* mature trophozoites were independently confirmed by RT-qPCR, and also observed at the late ring stage and in the knockout lines of D10 and HB3 genetic background (Supplementary Fig. 9c). These results indicate that PfAP2-HS contributes to regulating the basal expression of the same chaperone-encoding genes that it activates upon HS, among a few other genes. Together with the observation that the growth defect of the *pfap2-hs* knockout lines is attenuated at 35°C, this suggests that knockout parasites have reduced proteostasis capacity, such that at 37°C they are at the edge of proteostasis collapse. Parasite lines that can grow normally at 37°C in spite of PfAP2-HS deletions including the three AP2 domains therefore must have alternative pathways active to ensure basal proteostasis. We hypothesise that mutant parasites expressing truncated PfAP2-HS are frequently selected under culture conditions because the truncations don’t pose a fitness cost at 37°C in the lines in which they appear, and they prevent unnecessary activation of the HS response, which can be detrimental^29^, by unintended mild stress that may occur during culture.

Artemisinins are potent antimalarial drugs that kill parasites by causing general protein damage^12,13^, whereas resistance is associated with mutations in the Kelch13 protein^30,31^ and involves cellular stress response pathways such as the ubiquitin/proteasome system and the ER-based unfolded protein response (UPR)^12,13,32,33^. Since PfAP2-HS regulates the expression of key chaperones, we tested the sensitivity of PfAP2-HS-defficient lines to dihydroartemisinin (DHA), the active metabolite of artemisinins. In all four different genetic backgrounds (3D7, D10, HB3 and Gambian isolate), knockout of *pfap2-hs* (or truncation before D1) resulted in higher sensitivity to a pulse of DHA than in lines with full PfAP2-HS, both at the ring or the trophozoite stage, whereas 10G showed increased sensitivity only when exposed at the trophozoite stage (Fig. 3d, Supplementary Fig. 8h-i). These results indicate that deletion of PfAP2-HS renders parasites more sensitive to chemical proteotoxic stress, in addition to HS, likely as a consequence of basal defects in cellular proteostasis. We reasoned that if parasites lacking the PfAP2-HS protein bear constitutive proteome defects, they should have low tolerance to disruption of other factors involved in proteostasis maintenance. Indeed, the 10E_Δ*pfap2-hs* line was more sensitive to the proteasome inhibitor epoxomicin than the parental 10E line or the 10G line (Fig. 3e). Furthermore, after HS, there was more accumulation of polyubiquitinated proteins in the knockout line than in 10E or 10G, reflecting higher levels of unresolved protein damage (Supplementary Fig. 8j). We also assessed the links between the PfAP2-HS-driven HS response and the other main cell stress response pathway, the UPR. Using phosphorylation of eIF2α as a UPR marker, we found that the UPR does not depend on PfAP2-HS and is not directly activated by HS, because the marker was significantly elevated after HS only in the knockout line (Supplementary Fig. 8k).

Altogether, our results demonstrate that the PfAP2-HS transcription factor binds to the tandem G-box DNA motif at the promoter of a limited number of critical chaperone-encoding genes, *hsp70-1* and *hsp90*, and in response to febrile temperatures rapidly upregulates their expression (Fig. 3f). Specific binding to the G-box is conferred by D1^10^, but activation of these genes during HS strictly requires D3, which is not capable of binding DNA *in vitro*^10^ and likely participates in protein-protein interactions or dimerization^34^. Other components of the protein folding machinery necessary for HS survival^35-37^ are constitutively expressed or induced later, but the rapid PfAP2-HS-driven response is essential to avoid irreversible damage, as parasites lacking D3 or the full protein fail to survive HS. While PfAP2-HS is not evolutionarily related with HSF1^5,8^, the conserved master regulator of the HS response from yeast to mammals, it appears to play an analogous role. HSF1 also regulates a compact transcriptional program^6,7^ that includes the *hsp70* and *hsp90* genes. In yeast, the only essential role of this transcription factor is activating these two genes^7^. In addition to driving the protective HS response, PfAP2-HS also plays a D3-independent role under nonstress conditions, which is essential for growth at 37°C in some *P. falciparum* genetic backgrounds. Several lines of evidence indicate that the role of PfAP2-HS under basal conditions is related with proteostasis maintenance (Fig. 3f), as in the case of yeast HSF1^7^. Furthermore, the malarial HS response likely mediates protection against different types of proteotoxic stress, in addition to high temperature, as reported in other organisms^5,16^. This idea is supported by the observation that orthologs of *pfap2-hs* are present across *Plasmodium* spp. (Supplementary Fig. 10), including murine malaria species that do not induce fever. The identification of the PfAP2-HS transcription factor, which drives a strong protective response to an environmental condition, settles the controversy of whether malaria parasites can produce directed transcriptional responses to fluctuations in their within-host environment^38^ and provides a much-needed system to study induced transcriptional responses in malaria parasites.

## METHODS

### Parasite cultures

The 3D7-A stock of the clonal *P. falciparum* line 3D7^39^, the 3D7-A subclones 10G, 1.2B, 10E, 4D, 6D, 1.2F, W4-1, W4-2, W4-4, W4-5^40,41^, the HB3B (mosquito and chimpanzee-passaged HB3)^42^ and D10^43^ clonal parasite lines, and the culture-adapted Line 1 from The Gambia^26^ have been previously described. The HS-selected lines 3D7-A-HS r1 and r2 were derived from 3D7-A by exposing cultures to a 3 h HS for five consecutive cycles (each replicate, r1 and r2, is a fully independent selection from the 3D7-A stock), and the 3D7-A r1 and r2 lines are cultures maintained in parallel at 37°C without HS^19^. The generation of the new transgenic lines is described below. Parasites were cultured in B+ erythrocytes at a 3 % haematocrit under standard culture conditions in RPMI-based media containing Albumax II (without human serum), in a 5% CO_2_, 3% O_2_, balance N_2_ atmosphere (except for cultures for ChIP-seq experiments, in which O+ erythrocytes were used). Regular synchronization was performed using 5 % sorbitol lysis, whereas tight synchronization (1, 2 or 5 h age window) was achieved by Percoll purification followed by sorbitol treatment 1, 2 or 5 h later. All cultures were regularly maintained at 37°C, with the exception of the *pfap2-hs* knockout lines that were maintained at 35°C. For experiments performed in parallel with the knockout lines and other parasite lines, all cultures were maintained at 35 °C from at least one cycle before the experiment.

### Generation of transgenic parasite lines

Given the large size of *pfap2-hs* (11,577 bp), to knock out this gene with the CRISPR/Cas9 system we used two single guide RNAs (hereafter referred to as sgRNA or guide) (Supplementary Fig. 2a). One guide targets a sequence near the 5’ end of the gene (position 866-885 from the start codon) whereas the other recognizes a sequence near the 3’ end (positions 11,486-11,505). The 5’ guide was cloned into a modified pL6-*egfp* donor plasmid^44^ in which the *yfcu* cassette had been removed by digestion with *Not*I and *Sac*II, end blunting and re-ligation. 5’ and 3’ homology regions (HR1: positions -2 to 808 of the gene; HR2: positions +11,520 of the gene to 490 bp after the STOP codon) were also cloned in this plasmid, flanking the *hdhfr* expression cassette, to generate plasmid pL7-pfap2hs_KO_sgRNA5’. The 3’ guide was cloned into a modified version of the pDC2-Cas9-U6-hdhfr^45^ plasmid, in which we previously removed the *hdhfr* expression cassette by digesting with *Nco*I and *Sac*II, end blunting and re-ligation, and replaced the *Bbs*I guide cloning site by a *BtgZ*I site. The resulting plasmid was termed pDC2_wo/hdhfr_pfap2hs_sgRNA3’. All guides were cloned using the In-Fusion system (Takara) approximately as described^44^, whereas homology regions where PCR-amplified from genomic DNA and cloned by ligation using restriction sites *Spe*I and *Afl*II (HR1), and *EcoR*I and *Nco*I (HR2).

To tag PfAP2-HS with a 2xHA-ddFKBP domain tag (1.2B_ *pfap2hs*_ddFKBP line) we used a single homologous recombination approach (Supplementary Fig. 2b). To generate the pfap2hs_HA-ddFKBP plasmid, we replaced the *pfap2-g* homology region in plasmid PfAP2-G-ddFKBP^14^ by a PCR-amplified fragment including positions 9,551-11,574 of *pfap2-hs* in frame with the tag. The fragment was cloned using restriction sites *Not*I and *Xho*I.

For constructs aimed at C-terminal tagging of *pfap2-hs* using CRISPR/Cas9 (10E_*pfap2-hs*_eYFP-Cterm and 10E_*pfap2-hs*_3xHA-Cterm lines) we used a guide corresponding to positions 11,508-11,527 of the gene (Supplementary Fig. 2c-d). The guide was cloned in the pDC2-Cas9-U6-hdhfr^45^ plasmid to obtain pDC2_pfap2hs_sgRNA-C. The donor plasmid for tagging with eYFP (pHR-C_pfap2hs_eYFP) was based on plasmid pHRap2g-eYFP^46^, with the *pfap2-g* homology regions and *hsp90* 3’ sequence replaced by *pfap2-hs* homology regions. The 5’ homology region (HR1) was generated with a PCR-amplified fragment spanning from nucleotide 10,964 to the sequence of the guide (recodonized), and a 47 bp fragment (generated by annealing two complementary oligonucleotides) consisting of a recodonized version of the remaining nucleotides to the end of the gene. The two fragments were cloned simultaneously, using the In-Fusion system, into *Spe*I-*Bgl*II sites. The 3’ homology region (HR2) was a PCR fragment spanning position +1 to +590 after the *pfap2-hs* STOP codon. It was cloned into *Xho*I-*Aat*II restriction sites. The donor plasmid for 3xHA C-terminal tagging (pHR-C_pfap2hs_3xHA_hsp90-3’) was also based on plasmid pHRap2g-eYFP^46^, with the eYFP coding sequence replaced by the 3xHA sequence (amplified from plasmid pHH1inv-pfap2-g-HAx3^14^) and the same homology regions as in plasmid pHR-C_pfap2hs_eYFP (but HR2 was cloned, using the In-Fusion system, into *EcoR*I-*Aat*II sites, because in this construct the *hsp90* 3’ region in pHRap2g-eYFP was maintained).

For N-terminal tagging (10E_*pfap2-hs_*eYFP-Nterm line), we cloned a guide targeting *pfap2-hs* positions 73-92 in the pDC2-Cas9-U6-hDHFRyFCU^47^ plasmid to obtain plasmid pDC2_pfap2hs_sgRNA-N (Supplementary Fig. 2e). The donor plasmid (pfap2hs_HR-N_eYFP) consisted of a 5’ homology region (HR1) including positions -366 to -1 relative to the *pfap2-hs* start codon, the eYFP gene and an in frame 3’ homology region (HR2) spanning positions 4-756 of the gene (excluding the ATG) in which the nucleotides up to the position of the guide were recodonized. HR1 and HR2 were cloned into *Sac*II-*Nco*I and *Spe*I-*Eco*RI sites, respectively. HR2 was amplified in two steps using a nested PCR approach to add the recodonized sequences. The eYFP fragment (PCR-amplified from plasmid pHR-C_pfap2hs_eYFP) was cloned using the In-Fusion system into *Spe*I/*Nco*I sites. All oligonucleotides used to generate the plasmids are described in Supplementary Table 5.

Transfections were performed by electroporation of ring stage cultures with 100 µg of plasmid (HA-ddFKBP tagging) or with a mixture of 12 µg linearized donor plasmid and 60 µg of circular Cas9 plasmid (CRISPR-Cas9 system).

Linearization was achieved by digestion with the *Pvu*I restriction enzyme (cleaving the *amp* resistance gene of the donor plasmid). Transfected cultures were selected with 10 nM WR99210 for four days as previously described^47^ (transfections using the CRISPR-Cas9 system), or with continuous WR99210 pressure until parasites were observed, followed by 3 off/on drug cycles and subcloning by limiting dilution (transfections with the pfap2hs_HA-ddFKBP plasmid). In all cases, to assess correct integration we used analytical PCR of genomic DNA (Supplementary Fig. 2) with specific primers (Supplementary Table 5).

### HS resistance assay

HS was always performed on cultures at the mature trophozoite stage unless otherwise indicated. To measure survival to HS, cultures were tightly synchronized to a defined age window, diluted to 1% parasitaemia, split in two identical petri dishes (HS and control) maintained in independent air-tight incubation chambers, and exposed to HS when the majority of parasites were at the mature trophozoite stage (typically ∼30-35 h post-invasion, hpi; Δ*pfap2-hs* lines were tightly synchronized 3 h earlier than the other lines but exposed to HS in parallel to account for their slower IDC progression). The exception was experiments to screen many subclones (i.e., Fig. 1e) or to characterize transgenic parasite lines (i.e., Supplementary Fig. 2), in which cultures were only sorbitol-synchronized and HS performed ∼20-25 h after sorbitol lysis (mature trophozoite stage). For HS, the full incubation chamber was transferred to an incubator at 41.5°C for 3 h, and then placed back to 37 or 35°C (the latter temperature was used for all lines in experiments including the *pfap2-hs* knockout lines). The chamber with the control cultures was always maintained at 37 or 35°C. After reinvasion (typically ∼60-70 h after synchronization to ensure that all parasites had completed the cycle, including parasites subjected to HS that show delayed progression through the IDC), parasitaemia of control and HS-exposed cultures was measured by flow cytometry using a FACScalibur flow cytometer (Becton Dickinson) and SYTO 11 to stain nucleic acids, as previously described^48^.

### Phenotypic characterization

To determine the growth rate (increase in parasitaemia between consecutive cycles) at different temperatures, the parasitaemia of sorbitol-synchronized cultures was adjusted to 1% and then accurately determined by flow cytometry. Cultures where then split between two or three dishes maintained in parallel in incubators at the different temperatures tested. Parasitaemia was again determined by flow cytometry at the next cycle to determine the growth rate. To measure the duration of the IDC (at 35°C) in the different parasite lines we used a recently described method based on synchronization to a 1 h age window achieved by Percoll-purification of schizonts followed by sorbitol lysis 1 h later^48^. The determination of the number of merozoites per fully mature schizont was based on light microscopy analysis of Giemsa-stained smears from Percoll-purified schizonts^48^. DHA (Sigma no. D7439) or epoxomicin (Selleckchem no. S7038) sensitivity was measured by exposing tightly synchronized cultures at the ring (10-15 hpi, DHA only) or trophozoite (30-35 hpi, DHA or epoxomicin) stage (1% parasitaemia) to different concentrations of the compounds for 3 h. Parasitaemia was measured by light microscopy analysis of Giemsa-stained smears at the next cycle (typically 70-80 h after Percoll-sorbitol synchronization). For these experiments, the Δ*pfap2-hs* lines were tightly synchronized 3 h earlier than the other lines but exposed to DHA or epoxomicin in parallel (13-18 or 33-38 hpi), to account for their slower IDC progression.

### Transcriptional analysis by reverse transcription–quantitative PCR (RT-qPCR)

RNA from tightly synchronized cultures exposed to HS and their controls was obtained using the Trizol method, DNAse-treated and purified essentially as described^49^. Reverse transcription and qPCR analysis of cDNAs were also performed as described before^49^. In brief, a mixture of random primers and oligo (dT) were used for reverse transcription, and for qPCR we used the PowerSYBR Green Master Mix (Applied biosystems) and the standard curve method (each plate included a standard curve for each primer pair). All primers used are listed in Supplementary Table 5. Unless otherwise indicated, transcript levels were normalized against *serine--tRNA ligase* (PF3D7_0717700), which shows relatively stable expression throughout the IDC.

### Transcriptomic analysis using microarrays

To compare the transcriptome of control and HS-adapted 3D7-A parasite lines across the IDC we used previously described two-colour long oligonucleotide-based glass microarrays^19^. RNA was obtained from tightly synchronized cultures (5 h age window) at 8-13, 16-21, 24-29, 32-37 and 40-45 hpi. All samples (Cy5-labeled) were hybridized together with a reference pool (Cy3-labeled) consisting of a mixture of equal amounts of cDNA from rings, trophozoites and schizonts from control and HS-adapted lines. Comparative genome hybridization was used to identify potential transcript level differences attributable to genetic deletions or duplications. 5,142 genes produced useful data. Sample preparation, microarray hybridization and data analysis were performed essentially as described^19^.

To analyse the transcriptome of 10E, 10G and 10E_Δ*pfap2-hs* parasite lines under control and HS conditions, we used two-colour long oligonucleotide-based custom Agilent microarrays^50^. The microarray design was based on Agilent design AMADID no. 037237^50,51^, but we modified it as previously described (new design AMADID no. 084561)^52^. RNA was obtained from cultures synchronized to a 5 h age window at a ∼2.5% parasitaemia. Given the slower IDC progression of 10E_Δ*pfap2-hs*, cultures of this parasite line were synchronized to 0-5 hpi 3 h earlier than 10E and 10G cultures, such that at the time of starting HS (in parallel for all lines) all cultures were approximately at the same stage of IDC progression. HS was started at 30-35 hpi (33-38 hpi for the 10E_Δ*pfap2-hs* line) and samples collected before, during and after HS as indicated. RNA was prepared using the Trizol method. Sample preparation and microarray hybridization were performed essentially as described^51^. All samples (Cy5-labeled) were hybridized together with a reference pool (Cy3-labeled) consisting of a mixture of equal amounts of cDNA from rings, trophozoites and schizonts from 3D7-A. Microarray images were obtained using a DNA Microarray Scanner (no. G2505C, Agilent Technologies) located in a low ozone area, and initial data processing was performed using the GE2 _1105_Oct12 extraction protocol in Agilent Feature Extraction 11.5.1.1 software.

Agilent microarray data was analysed using Bioconductor in an R environment (R version 3.5.3). For each individual microarray, we calculated Cy3 and Cy5 background signal as the median of the 100 lowest signal probes for each channel, and probes with both Cy3 and Cy5 signals below three times the array background were excluded. Gene level log_2_(Cy5/Cy3) values, statistical estimation of parasite age^53^ and estimation of average expression fold-differences across a time interval (for the comparison between parasite lines in the absence of HS) were performed as described^19^. The log_2_ of the expression fold-change upon HS was calculated, for each gene and time point, as the log_2_(Cy5/Cy3) in the HS-exposed sample minus the log_2_(Cy5/Cy3) in the control sample at the same parasite age, calculated using linear interpolation in the log_2_(Cy5/Cy3) versus estimated age plot. Visual inspection was used to exclude from further analysis genes with apparent artefacts. Genes missing data for ≥2 time points (or ≥1 for the comparison between parasite lines in the absence of HS across a time interval), or with values within the lowest 15th percentile of expression intensity (Cy5 sample channel) in all samples, were also excluded from further analysis. 4,964 genes produced useful data.

To assess the level of similarity to a reference non-stressed transcriptome (HB3 line) with high temporal resolution^23^ we calculated the Pearson correlation between each sample and the time point with which it has higher similarity. Heatmaps and hierarchical clustering based on Spearman (Fig. 2) or Pearson (Supplementary Fig. 6) correlation were generated using Multiple Experiment Viewer (MeV) 4.9^54^. Expression trend plots for each cluster were generated using ggplot2, with LOESS smoothing, and Venn diagrams using the eulerr package (both in an R environment). Motif analysis (5 to 8 bp) was performed using MEME 5.0.3 software. Functional enrichment analysis using GO terms annotated in PlasmoDB version 43 was performed using Ontologizer 2.1 software^55^ with the topology-elim method^56^. Gene set enrichment analysis (GSEA) was performed using GSEA v3.0 Preranked^57^.

### Whole-genome sequencing analysis and analysis of publicly available genome sequences from field isolates

To sequence the full genome of control and HS-adapted 3D7-A lines (two biological replicates), we used PCR-free whole-genome Illumina sequencing. In brief, genomic DNA was sheared to ∼150-400 bp fragments using a Covaris S220 ultrasonicator and analysed using an Agilent 2100 Bioanalyzer. For library preparation we used the NEBNext DNA Library Prep Master Mix Set for Illumina (no. E6040S) using specific paired-end TruSeq Illumina adaptors for each sample. After quality check by qPCR, we obtained >6 million 150 bp paired reads for each sample using an Illumina MiSeq sequencing system. After checking reads quality (FastQC algorithm) and trimming adaptors (Cutadapt algorithm), sequence reads were mapped to the PlasmodDB *P. falciparum* 3D7 reference genome version 24 (https://plasmodb.org/plasmo/) using the Bowtie2 local alignment algorithm. Variant calling was performed using GATK-UnifiedGenotyper and following a pipeline based on GATK best practices to identify SNPs and small indels. Variants with low calling quality (Phred QUAL<20) and low read depth (DP<10) were filtered out using GATK-VariantFiltration, and only variants present in both biological replicates were considered. Differences in SNP/indel frequency between control and HS-adapted lines were calculated for each SNP/indel, and those showing <25% difference in any of the two replicates were filtered out. Genome Browse (Golden Helix) was used to visualize alignments and variants.

For the analysis of publicly available genome sequences, we used the Pf3K Project (2016): pilot data release 5 (www.malariagen.net/data/pf3k-5) containing the sequence of >2,500 field isolates. Only SNPs that passed all quality filters and did not fall within a region with multiple large insertions and deletions were considered to be high-confidence. Using these criteria, a single high-confidence polymorphism was identified at the *pfap2-hs* gene (producing the C3168X mutation that results in a truncated PfAP2-HS protein that lacks D3).

### ChIP experiments and data analysis

For ChIP experiments, synchronous 50 ml cultures at 2.5 to 5% parasitemia were harvested at the mid trophozoite stage. For replicates in which ChIP was performed in parallel under HS and control conditions, cultures were split off from a single parent flask at the mid trophozoite stage. Control flasks were immediately returned to 37°C whereas HS flasks were maintained at 41.5°C for 3 h before harvesting for ChIP analysis.

ChIP followed by qPCR or Illumina sequencing was performed as described^58^ using the 3F10 rat anti-HA antibody (1:500; Roche no. 11867423001) to immunoprecipitate HA tagged AP2-HS, with the following modification: total chromatin was diluted 5-fold in dilution buffer following sonication. The Illumina HiSeq system was used to obtain 125 bp paired-end (replicates 1-3) or 150 bp single-end (replicates 4-5) reads.

ChIP-seq data analysis was performed essentially as described^58^. In brief, after trimming, quality control, mapping the remaining reads to the *Plasmodium falciparum* genome (PlasmoDB release 28) using BWA-MEM and filtering duplicated reads, peak calling was performed using MACS2^59^ with a q-value cut-off of 0.01. Conversion to log_2_ coverage of immunoprecipitate/input was performed using DeepTools BamCompare, selecting the paired end parameter for all tools when analyzing experiments including control and HS conditions. Overlapping intervals within called peaks for each dataset were determined using Bedtools MultiIntersect. The closest annotated gene coding sequence for each called peak was determined using Bedtools ClosestBed. To visualize aligned data, we used IGV.

ChIP samples were analysed by qPCR in triplicate wells with primers described in Supplementary Table 5. All primer pairs were confirmed to have between 80 and 110% efficiency using sheared genomic DNA as a template control. The percent input was calculated using the formula 100*2^(Ct adjusted input – Ct IP)^.

### Western blot

Synchronized cultures at the mature trophozoite stage were exposed to a regular 3 h HS or to a 1.5 h DHA pulse (positive control for a condition known to produce proteotoxic stress and induce the UPR)^25,32^. Parasites were obtained using saponin lysis (0.15% w/v saponin) and pellets solubilized in 1x SDS-PAGE loading buffer with 4% β-mercaptoethanol and boiled at 95 °C for 5 min. Proteins were resolved by SDS-PAGE on 4-20% TGX Mini-PROTEAN gels (Bio-rad) and transferred to nitrocellulose membranes (Bio-rad). After blocking with 5% (w/v) bovine serum albumin (Biowest) in TBS-T (0.1% Tween 20 in tris buffered saline), membranes were incubated at 4°C overnight with the following primary antibodies: rabbit anti-ubiquitin (1:1000; Cell Signaling Technology no. 3933), rabbit anti-phospho-eIF2α (1:1000; Cell Signaling Technology no. 3398) and rabbit anti-histone H3 (1:1000; Cell Signaling Technology no. 9715). After incubation with a goat anti-rabbit IgG-peroxidase (1:5000; Millipore no. AP307P) secondary antibody, peroxidase was detected using the Pierce ECL Western Blotting Substrate (Thermo Fisher Scientific) in an ImageQuant LAS 4000 imaging system. To control for equal loading, parts of the membranes corresponding to different molecular weight ranges were separately hybridized with different antibodies. Signal quantification was performed using ImageJ.

### Statistical analysis

*P* values were calculated using a two-tailed *t*-test (equal variance). Only significant *P* values are shown in the figures. No statistical analysis was performed for experiments involving only two replicates. In all cases, *n* indicates independent biological replicates (i.e., samples were obtained from independent cultures). For cell cycle duration and DHA or epoxomicin survival experiments, values were fitted to a sigmoidal dose-response curve.

## Code availability

The scripts used for the analysis of microarray and next generation sequencing data will be made available upon reasonable request, without any restrictions.

## Data availability

The microarray data presented in Fig. 2 and Supplementary Fig. 1, 5, 6 and 9 has been deposited to the Gene Expression Omnibus (GEO) database with accession code GSE149394. Genome sequencing and ChIP-seq data presented in Fig. 1a, Fig. 2 and Supplementary Fig. 7 have been deposited to the Sequence Read Archive (SRA) database with accession codes PRJNA626524 and PRJNA670721, respectively. The authors declare that all other relevant data generated or analysed during this study are included in the Article or the Supplementary Information files. Materials and protocols are available from the corresponding author on reasonable request.

## ACKNOWLEDGMENTS

The authors thank J.J. López-Rubio (University of Montpellier) for plasmid pL6-*egfp*, M. Lee (Wellcome Sanger Institute) for plasmid pDC2-Cas9-U6-hdhfr and E. Knuepfer (The Francis Crick Institute) for plasmid pDC2-Cas9-U6-hDHFRyFCU. The authors also thank O. Llorà-Batlle and C. Sànchez-Guirado (ISGlobal) for assistance with the generation of plasmids used in this study, N. Rovira-Graells (ISGlobal) and A. Gupta. (Nanyang Technological University) for assistance with 3D7-A and 3D7-A-HS microarray experiments, O. Billker (Wellcome Sanger Institute) for experiments attempted in *P. berghei* and H. Ginsburg (The Hebrew University of Jerusalem) for providing data from the Malaria Parasite Metabolic Pathways. This publication uses data generated by the Pf3k project (www.malariagen.net/pf3k). This work was supported by grants from the Spanish Ministry of Economy and Competitiveness (MINECO)/ Agencia Estatal de Investigación (AEI) (SAF2013-43601-R, SAF2016-76190-R and PID2019-107232RB-I00 to A.C.), co-funded by the European Regional Development Fund (ERDF, European Union), and from NIH/NIAID (1R01 AI125565 to ML). E.T.-F. and L.M.-T. were supported by fellowships from the Spanish Ministry of Economy and Competitiveness (BES-2014-067901 and BES-2017-081079, respectively), co-funded by the European Social Fund (ESF). T.J.R. was supported by a training grant by NIH/NIGMS (T32 GM125592-01). This research is part of ISGlobal’s Program on the Molecular Mechanisms of Malaria, which is partially supported by the Fundación Ramón Areces. We acknowledge support from the Spanish Ministry of Science and Innovation through the “Centro de Excelencia Severo Ochoa 2019-2023” Program (CEX2018-000806-S), and support from the Generalitat de Catalunya through the CERCA Program.

## AUTHOR CONTRIBUTIONS

E.T.-F. performed all experiments except for those presented in Supplementary Fig. 1, Western blot and ChIP-seq experiments. L.M.-T., E.T.-F., T.J.R. and A.C. performed the bioinformatics analysis. N.C.-V. performed Western blot experiments. T.J.R. performed and M.L. supervised ChIP-seq experiments. Z.B. provided microarray hybridizations for experiments presented in Supplementary Fig. 1. D.J.C. advised on clinical isolates and provided Line 1 from The Gambia. E.T.-F. and A.C. conceived the project, designed and interpreted the experiments, and wrote the manuscript (with input from all authors and major input from M.L. and D.J.C.).

