## Supplementary Figures for "The PfAP2-HS transcription factor protects malaria parasites from febrile temperatures"

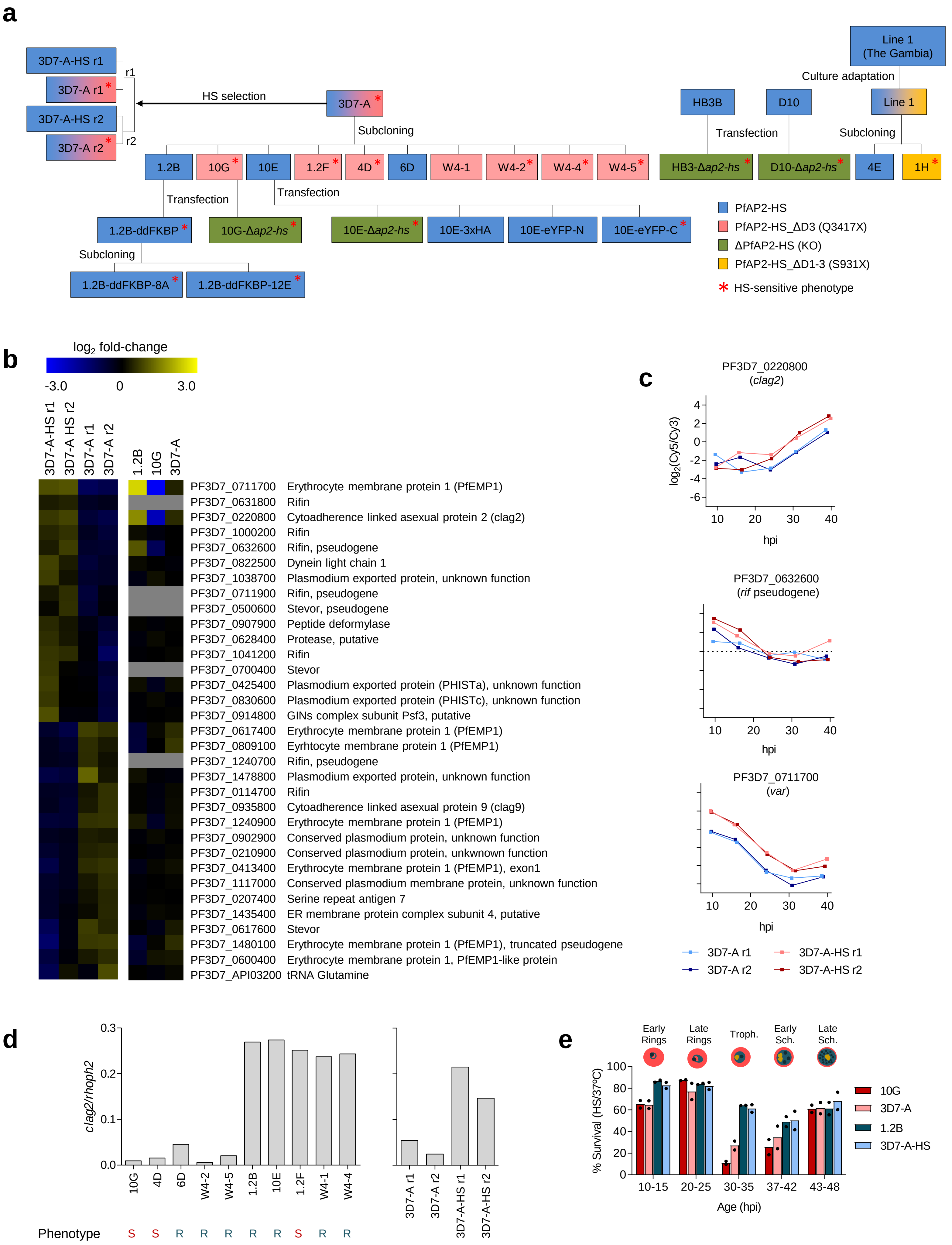

a
3D7-A-HS r1
3D7-A r1
r1
3D7-A-HS r2
3D7-A r2
r2
HS selection
3D7-A
Subcloning
1.2B
10G
10E
1.2F
4D
6D
W4-1
W4-2
W4-4
W4-5
Transfection
Transfection
1.2B-ddFKBP
10G-Δap2-hs
10E-Δap2-hs
10E-3xHA
10E-eYFP-N
10E-eYFP-C
Subcloning
1.2B-ddFKBP-8A
1.2B-ddFKBP-12E
Line 1
(The Gambia)
Culture adaptation
HB3B
D10
Line 1
Transfection
Subcloning
HB3-Δap2-hs
D10-Δap2-hs
4E
1H
PfAP2-HS
PfAP2-HS_ΔD3 (Q3417X)
ΔPfAP2-HS (KO)
PfAP2-HS_ΔD1-3 (S931X)
* HS-sensitive phenotype
b
log2 fold-change
-3.0
0
3.0
3D7-A-HS r1
3D7-A HS r2
3D7-A r1
3D7-A r2
1.2B
10G
3D7-A
PF3D7_0711700 Erythrocyte membrane protein 1 (PfEMP1)
PF3D7_0631800 Rifin
PF3D7_0220800 Cytoadherence linked asexual protein 2 (clag2)
PF3D7_1000200 Rifin
PF3D7_0632600 Rifin, pseudogene
PF3D7_0822500 Dynein light chain 1
PF3D7_1038700 Plasmodium exported protein, unknown function
PF3D7_0711900 Rifin, pseudogene
PF3D7_0500600 Stevor, pseudogene
PF3D7_0907900 Peptide deformylase
PF3D7_0628400 Protease, putative
PF3D7_1041200 Rifin
PF3D7_0700400 Stevor
PF3D7_0425400 Plasmodium exported protein (PHISTa), unknown function
PF3D7_0830600 Plasmodium exported protein (PHISTc), unknown function
PF3D7_0914800 GINs complex subunit Psf3, putative
PF3D7_0617400 Erythrocyte membrane protein 1 (PfEMP1)
PF3D7_0809100 Eyrhtocyte membrane protein 1 (PfEMP1)
PF3D7_1240700 Rifin, pseudogene
PF3D7_1478800 Plasmodium exported protein, unknown function
PF3D7_0114700 Rifin
PF3D7_0935800 Cytoadherence linked asexual protein 9 (clag9)
PF3D7_1240900 Erythrocyte membrane protein 1 (PfEMP1)
PF3D7_0902900 Conserved plasmodium protein, unknown function
PF3D7_0210900 Conserved plasmodium protein, unkwnown function
PF3D7_0413400 Erythrocyte membrane protein 1 (PfEMP1), exon1
PF3D7_1117000 Conserved plasmodium membrane protein, unknown function
PF3D7_0207400 Serine repeat antigen 7
PF3D7_1435400 ER membrane protein complex subunit 4, putative
PF3D7_0617600 Stevor
PF3D7_1480100 Erythrocyte membrane protein 1 (PfEMP1), truncated pseudogene
PF3D7_0600400 Erythrocyte membrane protein 1, PfEMP1-like protein
PF3D7_API03200 tRNA Glutamine
c
PF3D7_0220800
(clag2)
log2(Cy5/Cy3)
4
2
0
-2
-4
-6
10
20
30
40
hpi
PF3D7_0632600
(rif pseudogene)
10
20
30
40
hpi
PF3D7_0711700
(var)
10
20
30
40
hpi
3D7-A r1
3D7-A r2
3D7-A-HS r1
3D7-A-HS r2
d
clag2/rhoph2
0.3
0.2
0.1
0.0
10G
4D
6D
W4-2
W4-5
1.2B
10E
1.2F
W4-1
W4-4
3D7-A r1
3D7-A r2
3D7-A-HS r1
3D7-A-HS r2
Phenotype
S S R R R R R S R R
e
Early Rings
Late Rings
Troph.
Early Sch.
Late Sch.
% Survival (HS/37ºC)
100
80
60
40
20
0
10-15
20-25
30-35
37-42
43-48
Age (hpi)
10G
3D7-A
1.2B
3D7-A-HS

**Supplementary Figure 1. Transcriptional analysis of HS-adapted and control lines. a**, Schematic of the parasite lines used in this study. The colour code indicates whether they express wild type PfAP2-HS or truncated forms lacking AP2 domain 3, the three AP2 domains, or virtually the full protein (KO). Parasite lines shown with a colour gradient consist of a mixture of individual parasites expressing different versions of the protein. An asterisk indicates a HS sensitive phenotype, and r1 and r2 are independent replicates of the HS selection of 3D7-A over five consecutive cycles (3D7-A-HS r1 and r2 are the selected lines, whereas 3D7-A r1 and r2 are controls maintained in parallel at 37ºC during each selection). Mutations in *pfap2-hs* arose spontaneously in 3D7-A, and are not known to occur in other stocks of 3D7 (e.g., they don't occur in the reference 3D7 genome available in PlasmoDB). **b,** Microarray comparison of HS-adapted and control parasite lines across the asexual blood cycle. Values are the $\log_2$ of the maximum expression fold-change (from the average of all lines compared) across a time interval corresponding to half the length of the asexual cycle, calculated using the aMAFC score as previously described[1]. Genes with a >1.5-fold-change in expression in two independent 3D7-A HS-adapted lines (3D7-A-HS r1 and r2) relative to their respective controls (3D7-A r1 and r2) are shown. Data for parasite lines 10G (HS-sensitive subclone), 1.2B (HS-resistant subclone) and 3D7-A (right panel) is from Rovira-Graells et al.[1]. **c**, Time-course expression of genes in panel **b** that showed a concordant change in expression between HS-adapted and control cultures, and between the HS-resistant subclone 1.2B and the HS-sensitive subclone 10G. Based on the predicted function of the three genes, *clag2* was considered the most plausible candidate to play a role in HS resistance. **d**, Expression of *clag2* is neither necessary nor sufficient for HS resistance. RT-qPCR analysis of *clag2* transcript levels (normalized against *rhoph2*) in schizonts of HS sensitive (S) and HS resistant (R) 3D7-A subclones (see Fig. 1e), and of the HS-adapted and control lines. **e**, Survival of tightly synchronized cultures exposed to HS at different ages (in h post-invasion, hpi) for two HS-sensitive (3D7-A and 10G) and two HS-resistant (3D7-A-HS and 1.2B) lines (mean of $n$=2).

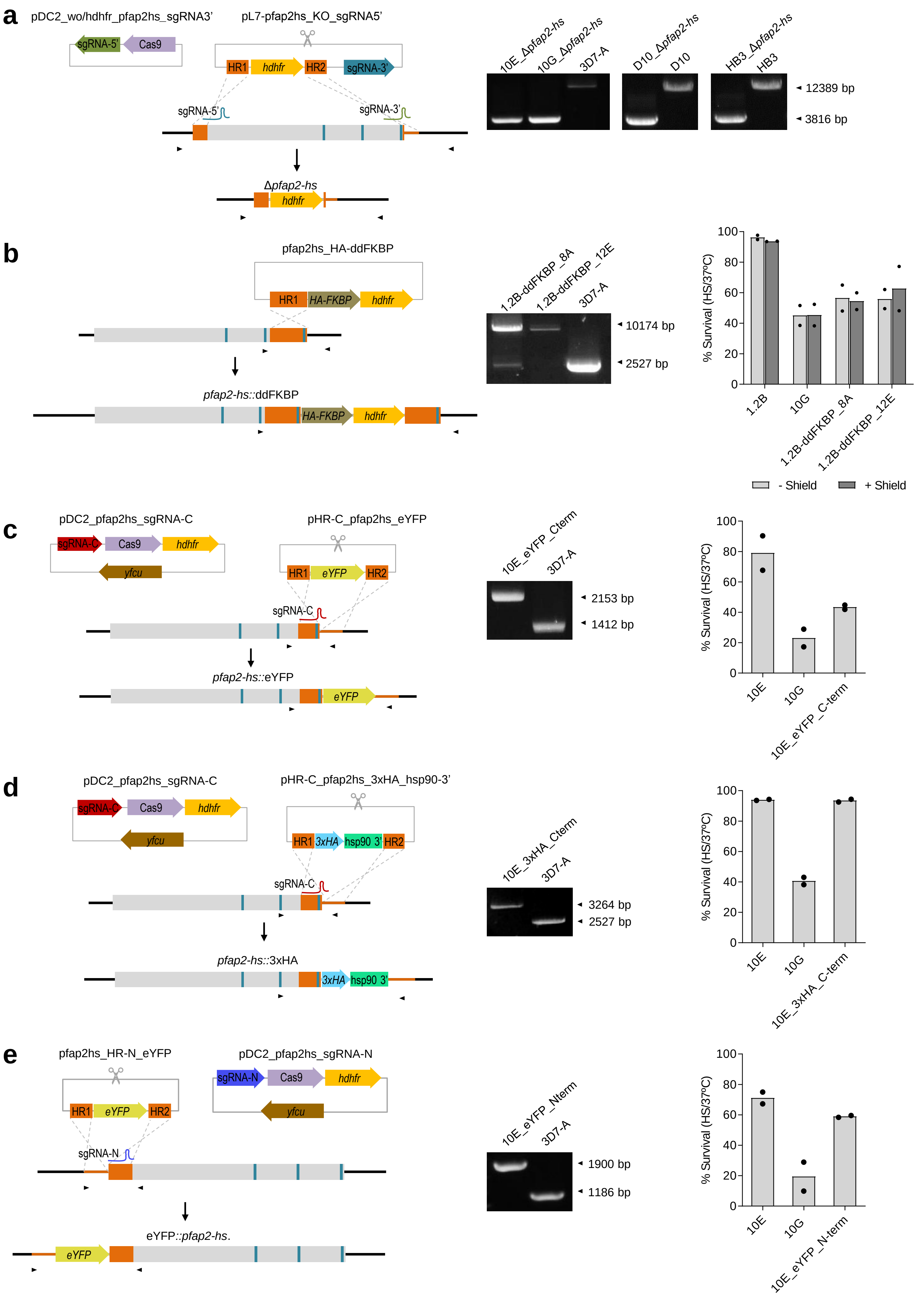

a
pDC2_wo/hdhfr_pfap2hs_sgRNA3'
sgRNA-5'
Cas9
pL7-pfap2hs_KO_sgRNA5'
HR1
hdhfr
HR2
sgRNA-3'
sgRNA-5'
sgRNA-3'
Δpfap2-hs
hdhfr
10E_Δpfap2-hs
10G_Δpfap2-hs
3D7-A
D10_Δpfap2-hs
D10
HB3_Δpfap2-hs
HB3
12389 bp
3816 bp
b
pfap2hs_HA-ddFKBP
HR1
HA-FKBP
hdhfr
pfap2-hs::ddFKBP
HA-FKBP
hdhfr
1.2B-ddFKBP_8A
1.2B-ddFKBP_12E
3D7-A
10174 bp
2527 bp
% Survival (HS/37ºC)
1.2B
10G
1.2B-ddFKBP_8A
1.2B-ddFKBP_12E
- Shield
+ Shield
c
pDC2_pfap2hs_sgRNA-C
sgRNA-C
Cas9
hdhfr
yfcu
pHR-C_pfap2hs_eYFP
HR1
eYFP
HR2
sgRNA-C
pfap2-hs::eYFP
eYFP
10E_eYFP_Cterm
3D7-A
2153 bp
1412 bp
% Survival (HS/37ºC)
10E
10G
10E_eYFP_C-term
d
pDC2_pfap2hs_sgRNA-C
sgRNA-C
Cas9
hdhfr
yfcu
pHR-C_pfap2hs_3xHA_hsp90-3'
HR1
3xHA
hsp90 3'
HR2
sgRNA-C
pfap2-hs::3xHA
3xHA
hsp90 3'
10E_3xHA_Cterm
3D7-A
3264 bp
2527 bp
% Survival (HS/37ºC)
10E
10G
10E_3xHA_C-term
e
pfap2hs_HR-N_eYFP
HR1
eYFP
HR2
pDC2_pfap2hs_sgRNA-N
sgRNA-N
Cas9
hdhfr
yfcu
sgRNA-N
eYFP::pfap2-hs.
eYFP
10E_eYFP_Nterm
3D7-A
1900 bp
1186 bp
% Survival (HS/37ºC)
10E
10G
10E_eYFP_N-term

**Supplementary Figure 2. Generation and characterization of transgenic parasite lines edited at the *pfap2-hs* locus. a**, Schematic of the CRISPR/Cas9 strategy used to knockout *pfap2-hs*, using two guide RNAs. **b**, C-terminal tagging of endogenous PfAP2-HS by single homologous recombination with a tag consisting of a 2xHA epitope and an FKBP destabilization domain (DD domain). **c-e**, Tagging of endogenous PfAP2-HS using CRISPR-Cas9 technology. The tags used where a C-terminal eYFP (**c**), a C-terminal 3xHA (**d**) and an N-terminal eYFP (**e**). In all panels, the position of the primers used for analytical PCR (arrowheads), guide RNA and AP2 domains (blue vertical bars) is indicated. The electrophoresis images at the right are the analytical PCR validation of the genetic edition, showing correct edition and absence of wild-type locus in all cases except for the 8A subclone of 1.2B-ddFKBP (8A and 12E are subclones obtained after drug cycling). The bar charts at the right show the level of survival (mean of *n*=2) of the transgenic lines upon HS, with HS-resistant (10E) and HS-sensitive (10G, expressing PfAP2-HSΔD3) subclones as controls. Addition of a C-terminal eYFP or HA-FKBP tag did not affect growth at 37ºC but resulted in high HS sensitivity, similar to the 10G line. In contrast, C-terminal addition of the smaller 3xHA tag or addition of an N-terminal eYFP did not affect growth at 37ºC or HS sensitivity. In all cases, tagged PfAP2-HS was not detectable by immunofluorescence or Western blot analysis, probably as a consequence of its very low abundance (see proteomic data in www.PlasmoDB.org).

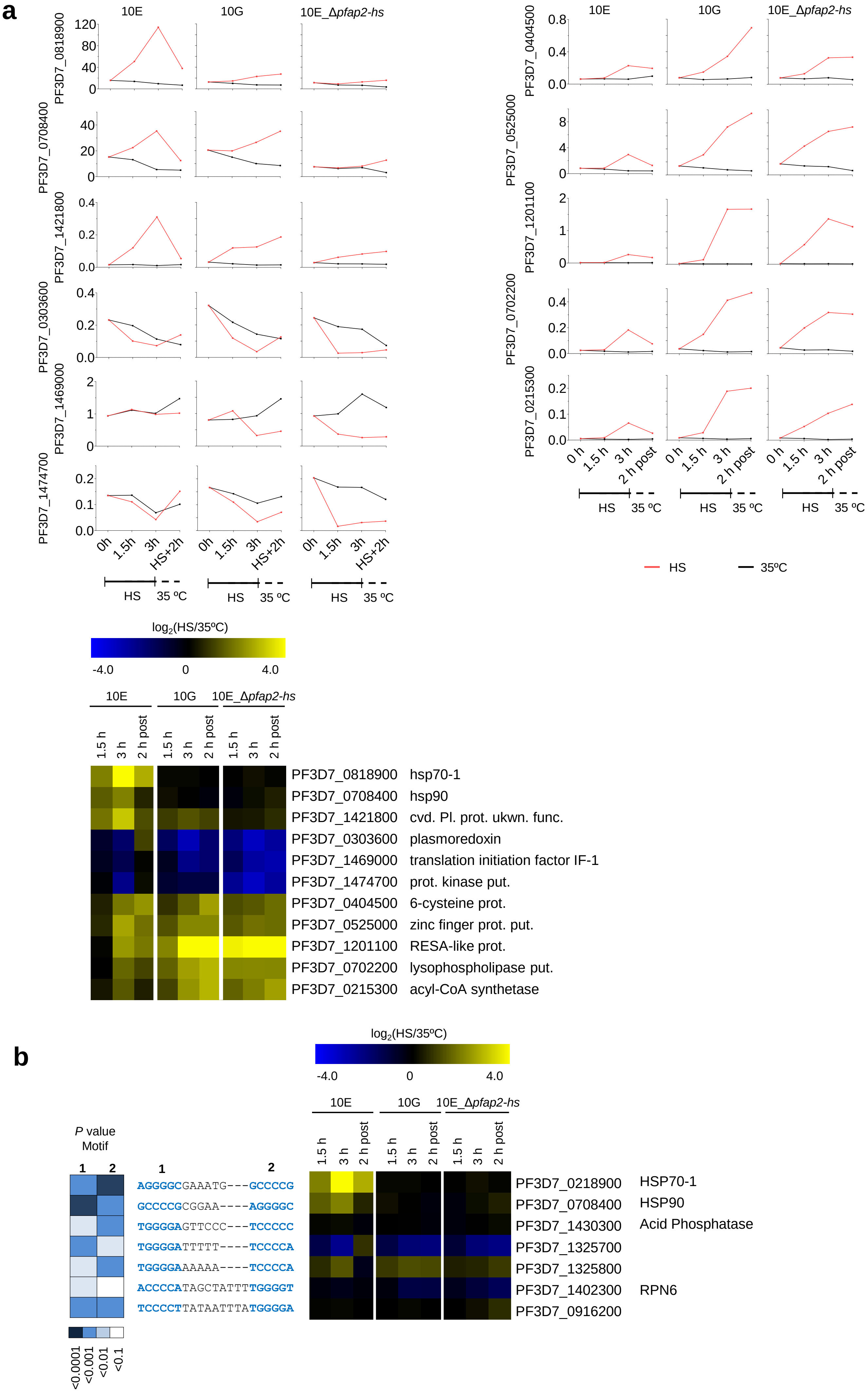
a
10E
10G
10E_Δpfap2-hs
PF3D7_0818900
PF3D7_0708400
PF3D7_1421800
PF3D7_0303600
PF3D7_1469000
PF3D7_1474700
PF3D7_0404500
PF3D7_0525000
PF3D7_1201100
PF3D7_0702200
PF3D7_0215300
0h
1.5h
3h
HS+2h
0 h
1.5 h
3 h
2 h post
HS
35 ºC
HS
35ºC
log2(HS/35ºC)
-4.0
0
4.0
1.5 h
3 h
2 h post
PF3D7_0818900 hsp70-1
PF3D7_0708400 hsp90
PF3D7_1421800 cvd. Pl. prot. ukwn. func.
PF3D7_0303600 plasmoredoxin
PF3D7_1469000 translation initiation factor IF-1
PF3D7_1474700 prot. kinase put.
PF3D7_0404500 6-cysteine prot.
PF3D7_0525000 zinc finger prot. put.
PF3D7_1201100 RESA-like prot.
PF3D7_0702200 lysophospholipase put.
PF3D7_0215300 acyl-CoA synthetase
b
P value
Motif
1
2
AGGGGCGAAATG---GCCCCG
GCCCCGCGGAA----AGGGGC
TGGGGAGTTCCC---TCCCCC
TGGGGATTTTT----TCCCCA
TGGGGAAAAAA----TCCCCA
ACCCCATAGCTATTTTGGGGT
TCCCCTTATAATTTATGGGGA
<0.0001
<0.001
<0.01
<0.1
PF3D7_0218900 HSP70-1
PF3D7_0708400 HSP90
PF3D7_1430300 Acid Phosphatase
PF3D7_1325700
PF3D7_1325800
PF3D7_1402300 RPN6
PF3D7_0916200

**Supplementary Figure 3. Validation of the transcriptomic changes upon HS and distribution of the tandem G-box motif. a**, RT-qPCR analysis of transcript levels (normalized against *serine--tRNA ligase*) of the genes selected for validation, using biological samples independent from the samples used for microarray analysis. Values are the average of triplicate reactions. The $log_2$ expression fold-change (HS relative to control conditions) for these genes in the microarray analysis (Fig. 2a) is shown to facilitate comparison. **b**, Genes in the *P. falciparum* genome containing tandem arrangements (maximum distance between the two: 9 nucleotides) of the G-box [(A/G)NGGGG(C/A)] motif in their regulatory regions (defined as the 2 kb upstream of the start codon or until the neighbour gene, when it is closer). The sequence of the G-box in each gene is shown in blue, and the level of concordance with the consensus G-box motif is expressed as the *P* value of the match (determined using the FIMO v5.0.5 function in the MEME suite). Expression changes upon HS for these genes are shown as in panel **a**.

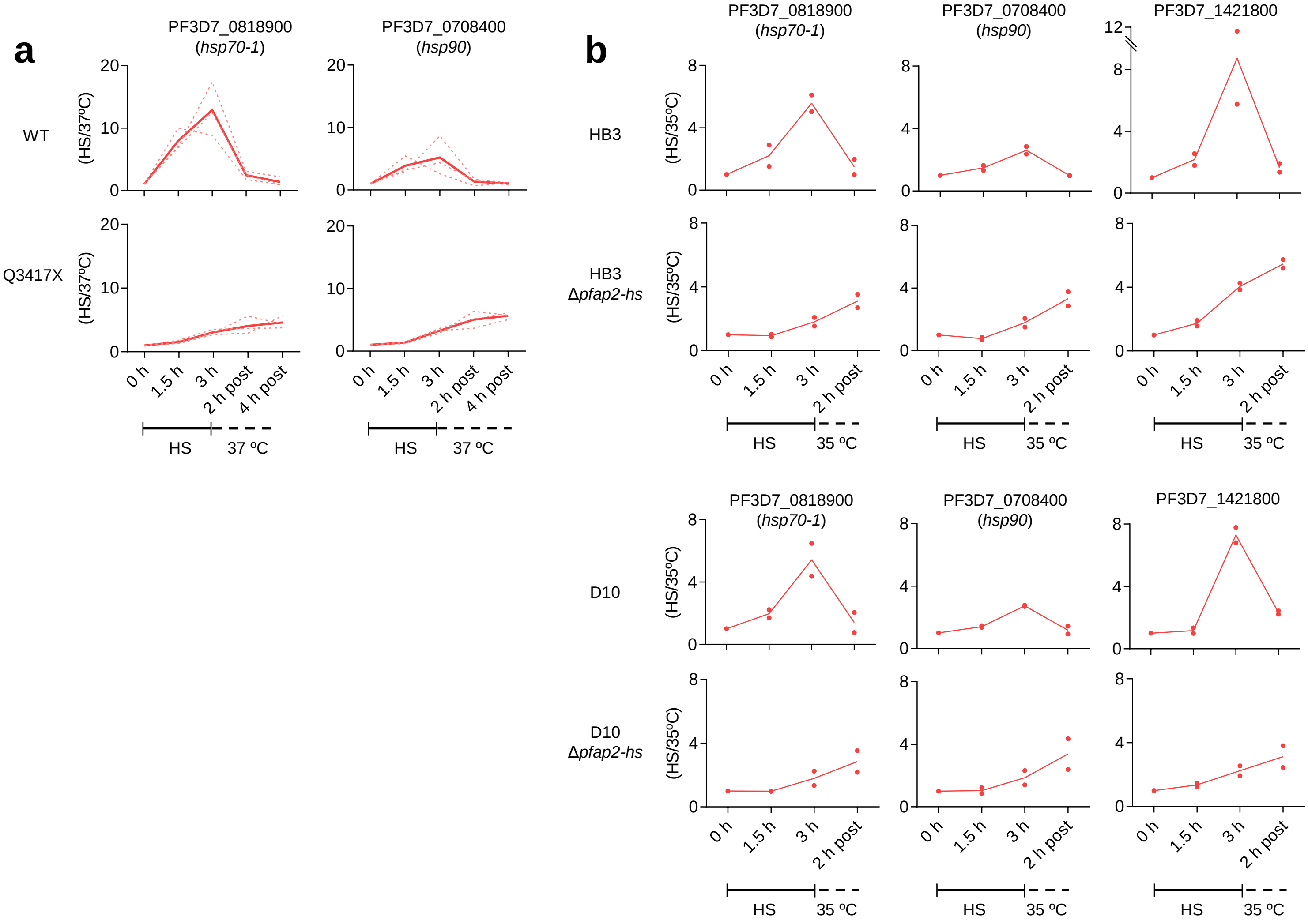


**Supplementary Figure 4. Changes in *hsp70-1, hsp90* and PF3D7_1421800 transcript levels in parasites lacking complete PfAP2-HS or D3.** Fold-increase in transcript levels (determined by RT-qPCR, normalized against *serine--tRNA ligase*) during and after HS starting at 33-35 (**a**) or 30-35 (**b**) hpi, relative to cultures maintained in parallel without HS. In panel **a**, values for three individual 3D7-A subclones carrying or not the Q3417X mutation are shown as dotted lines, whereas the average of the three subclones is shown as a continuous line. In panel **b**, the mean of *n*=2 independent biological replicates is shown.

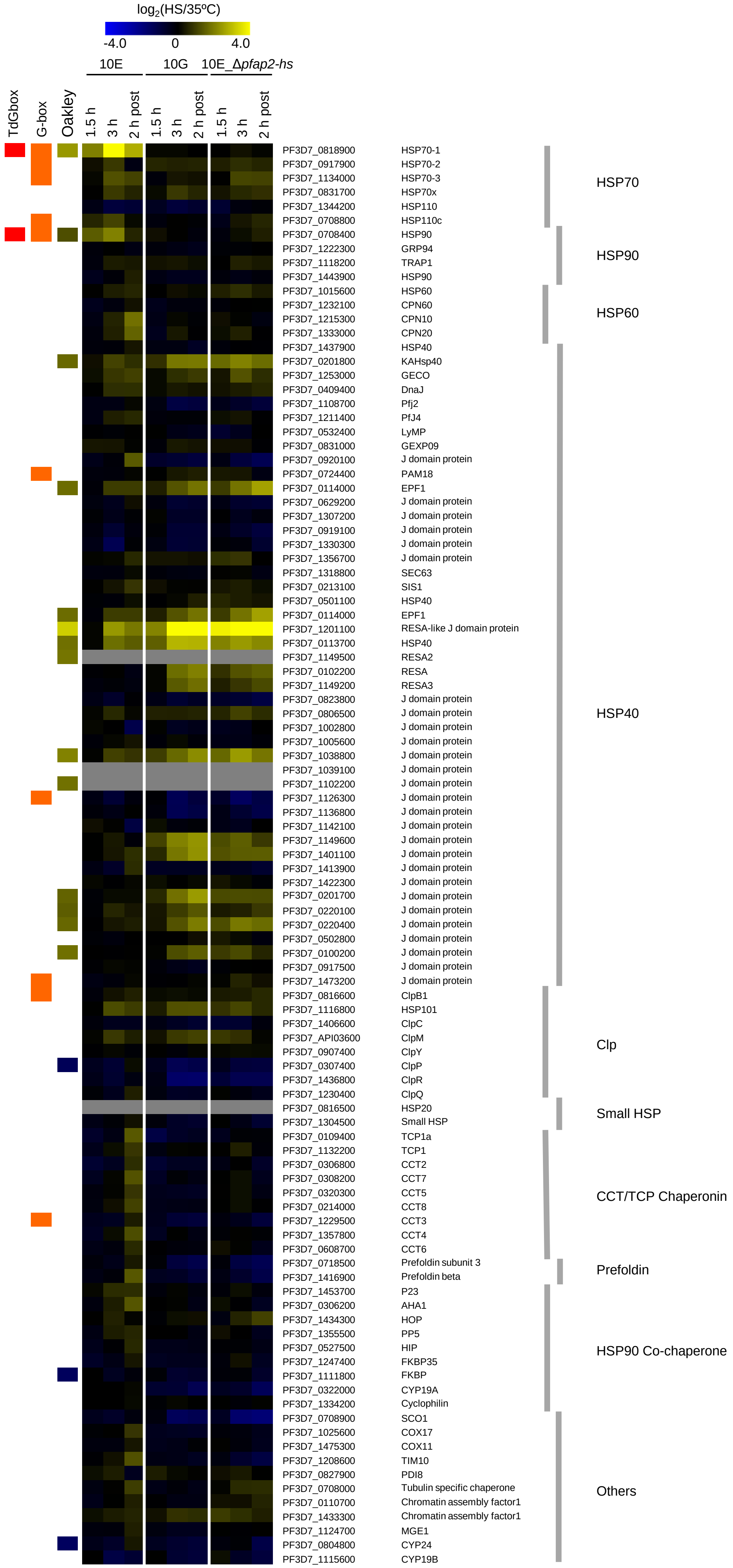

log2(HS/35ºC)
-4.0
0
4.0
10E
10G
10E_Δpfap2-hs
TdGbox
G-box
Oakley
1.5 h
3 h
2 h post
1.5 h
3 h
2 h post
1.5 h
3 h
2 h post
PF3D7_0818900 HSP70-1
PF3D7_0917900 HSP70-2
PF3D7_1134000 HSP70-3
PF3D7_0831700 HSP70x
PF3D7_1344200 HSP110
PF3D7_0708800 HSP110c
PF3D7_0708400 HSP90
PF3D7_1222300 GRP94
PF3D7_1118200 TRAP1
PF3D7_1443900 HSP90
PF3D7_1015600 HSP60
PF3D7_1232100 CPN60
PF3D7_1215300 CPN10
PF3D7_1333000 CPN20
PF3D7_1437900 HSP40
PF3D7_0201800 KAHsp40
PF3D7_1253000 GECO
PF3D7_0409400 DnaJ
PF3D7_1108700 Pfj2
PF3D7_1211400 PfJ4
PF3D7_0532400 LyMP
PF3D7_0831000 GEXP09
PF3D7_0920100 J domain protein
PF3D7_0724400 PAM18
PF3D7_0114000 EPF1
PF3D7_0629200 J domain protein
PF3D7_1307200 J domain protein
PF3D7_0919100 J domain protein
PF3D7_1330300 J domain protein
PF3D7_1356700 J domain protein
PF3D7_1318800 SEC63
PF3D7_0213100 SIS1
PF3D7_0501100 HSP40
PF3D7_0114000 EPF1
PF3D7_1201100 RESA-like J domain protein
PF3D7_0113700 HSP40
PF3D7_1149500 RESA2
PF3D7_0102200 RESA
PF3D7_1149200 RESA3
PF3D7_0823800 J domain protein
PF3D7_0806500 J domain protein
PF3D7_1002800 J domain protein
PF3D7_1005600 J domain protein
PF3D7_1038800 J domain protein
PF3D7_1039100 J domain protein
PF3D7_1102200 J domain protein
PF3D7_1126300 J domain protein
PF3D7_1136800 J domain protein
PF3D7_1142100 J domain protein
PF3D7_1149600 J domain protein
PF3D7_1401100 J domain protein
PF3D7_1413900 J domain protein
PF3D7_1422300 J domain protein
PF3D7_0201700 J domain protein
PF3D7_0220100 J domain protein
PF3D7_0220400 J domain protein
PF3D7_0502800 J domain protein
PF3D7_0100200 J domain protein
PF3D7_0917500 J domain protein
PF3D7_1473200 J domain protein
PF3D7_0816600 ClpB1
PF3D7_1116800 HSP101
PF3D7_1406600 ClpC
PF3D7_API03600 ClpM
PF3D7_0907400 ClpY
PF3D7_0307400 ClpP
PF3D7_1436800 ClpR
PF3D7_1230400 ClpQ
PF3D7_0816500 HSP20
PF3D7_1304500 Small HSP
PF3D7_0109400 TCP1a
PF3D7_1132200 TCP1
PF3D7_0306800 CCT2
PF3D7_0308200 CCT7
PF3D7_0320300 CCT5
PF3D7_0214000 CCT8
PF3D7_1229500 CCT3
PF3D7_1357800 CCT4
PF3D7_0608700 CCT6
PF3D7_0718500 Prefoldin subunit 3
PF3D7_1416900 Prefoldin beta
PF3D7_1453700 P23
PF3D7_0306200 AHA1
PF3D7_1434300 HOP
PF3D7_1355500 PP5
PF3D7_0527500 HIP
PF3D7_1247400 FKBP35
PF3D7_1111800 FKBP
PF3D7_0322000 CYP19A
PF3D7_1334200 Cyclophilin
PF3D7_0708900 SCO1
PF3D7_1025600 COX17
PF3D7_1475300 COX11
PF3D7_1208600 TIM10
PF3D7_0827900 PDI8
PF3D7_0708000 Tubulin specific chaperone
PF3D7_0110700 Chromatin assembly factor1
PF3D7_1433300 Chromatin assembly factor1
PF3D7_1124700 MGE1
PF3D7_0804800 CYP24
PF3D7_1115600 CYP19B
HSP70
HSP90
HSP60
HSP40
Clp
Small HSP
CCT/TCP Chaperonin
Prefoldin
HSP90 Co-chaperone
Others

**Supplementary Figure 5. Transcript level changes upon HS in chaperone-encoding genes.** $Log_2$ expression fold-change (HS relative to control conditions, as in Fig. 2a) for all chaperone-encoding genes described by Pavithra and colleagues[2]. Columns at the left indicate presence of the G-box[3] or tandem G-box (TdGbox) in the upstream region, and $log_2$ fold-change during HS in a previous study[4] (Oakley).

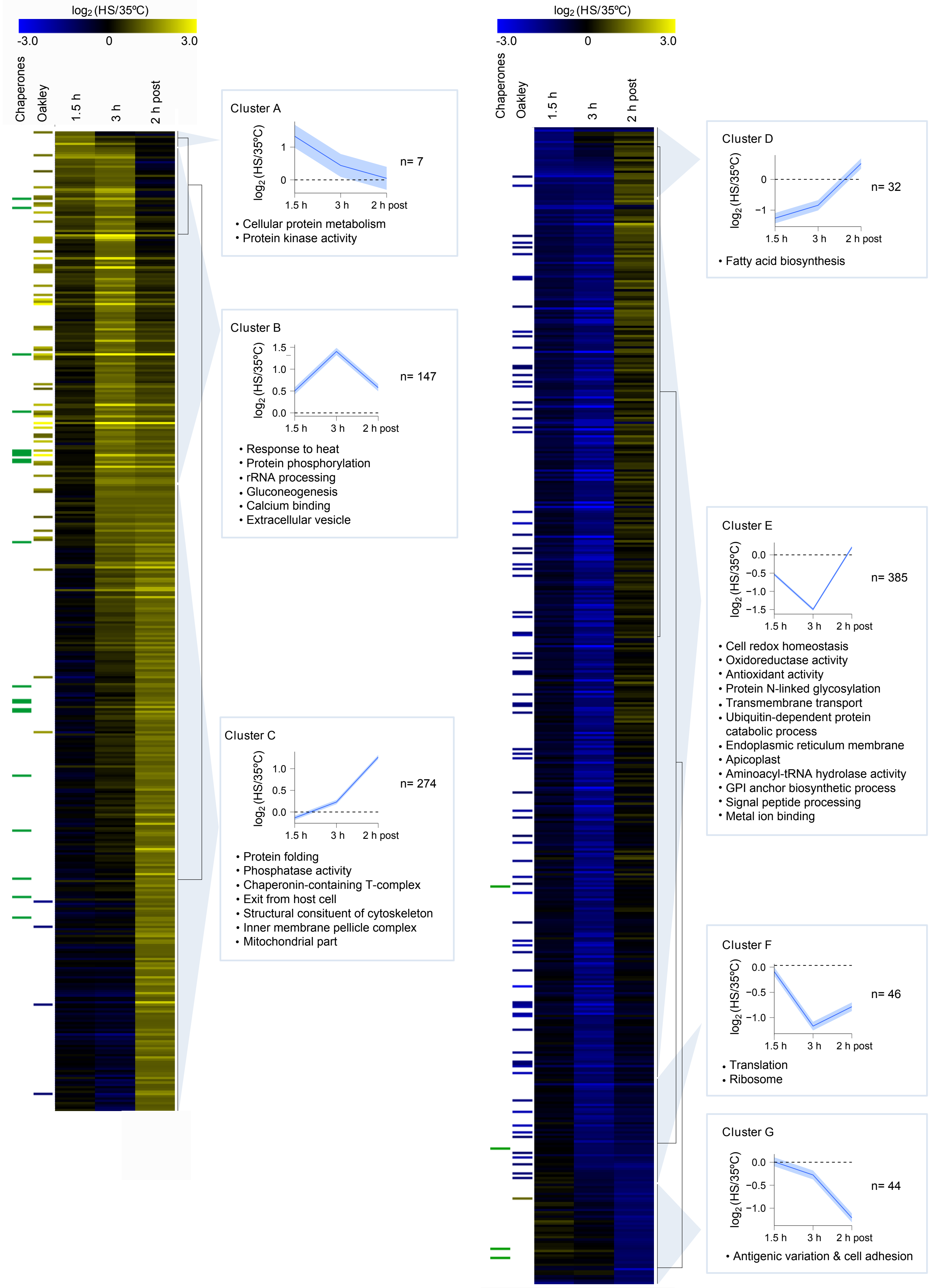

log2 (HS/35ºC)
-3.0
0
3.0
Chaperones
Oakley
1.5 h
3 h
2 h post
Cluster A
log2 (HS/35ºC)
n= 7
1.5 h
3 h
2 h post
• Cellular protein metabolism
• Protein kinase activity
Cluster B
log2 (HS/35ºC)
n= 147
• Response to heat
• Protein phosphorylation
• rRNA processing
• Gluconeogenesis
• Calcium binding
• Extracellular vesicle
Cluster C
log2 (HS/35ºC)
n= 274
• Protein folding
• Phosphatase activity
• Chaperonin-containing T-complex
• Exit from host cell
• Structural consituent of cytoskeleton
• Inner membrane pellicle complex
• Mitochondrial part
Cluster D
log2 (HS/35ºC)
n= 32
• Fatty acid biosynthesis
Cluster E
log2 (HS/35ºC)
n= 385
• Cell redox homeostasis
• Oxidoreductase activity
• Antioxidant activity
• Protein N-linked glycosylation
• Transmembrane transport
• Ubiquitin-dependent protein catabolic process
• Endoplasmic reticulum membrane
• Apicoplast
• Aminoacyl-tRNA hydrolase activity
• GPI anchor biosynthetic process
• Signal peptide processing
• Metal ion binding
Cluster F
log2 (HS/35ºC)
n= 46
• Translation
• Ribosome
Cluster G
log2 (HS/35ºC)
n= 44
• Antigenic variation & cell adhesion

**Supplementary Figure 6. Transcriptomic characterization of the HS response in parasites expressing complete PfAP2-HS (10E line).** $Log_2$ expression fold-change (HS relative to control conditions) in the wild-type 10E line determined by microarray analysis. Only genes with a fold-change ≥2 at any of the time points analysed are shown. The mean $log_2$ expression fold-change (with 95% confidence interval) and representative enriched GO terms are shown for each cluster. Columns at the left indicate fold-change during HS in a previous study[4] (Oakley), and annotation as chaperone[2]. Ten genes had values out of the range displayed (actual range: -3.89 to +4.03).

**a**

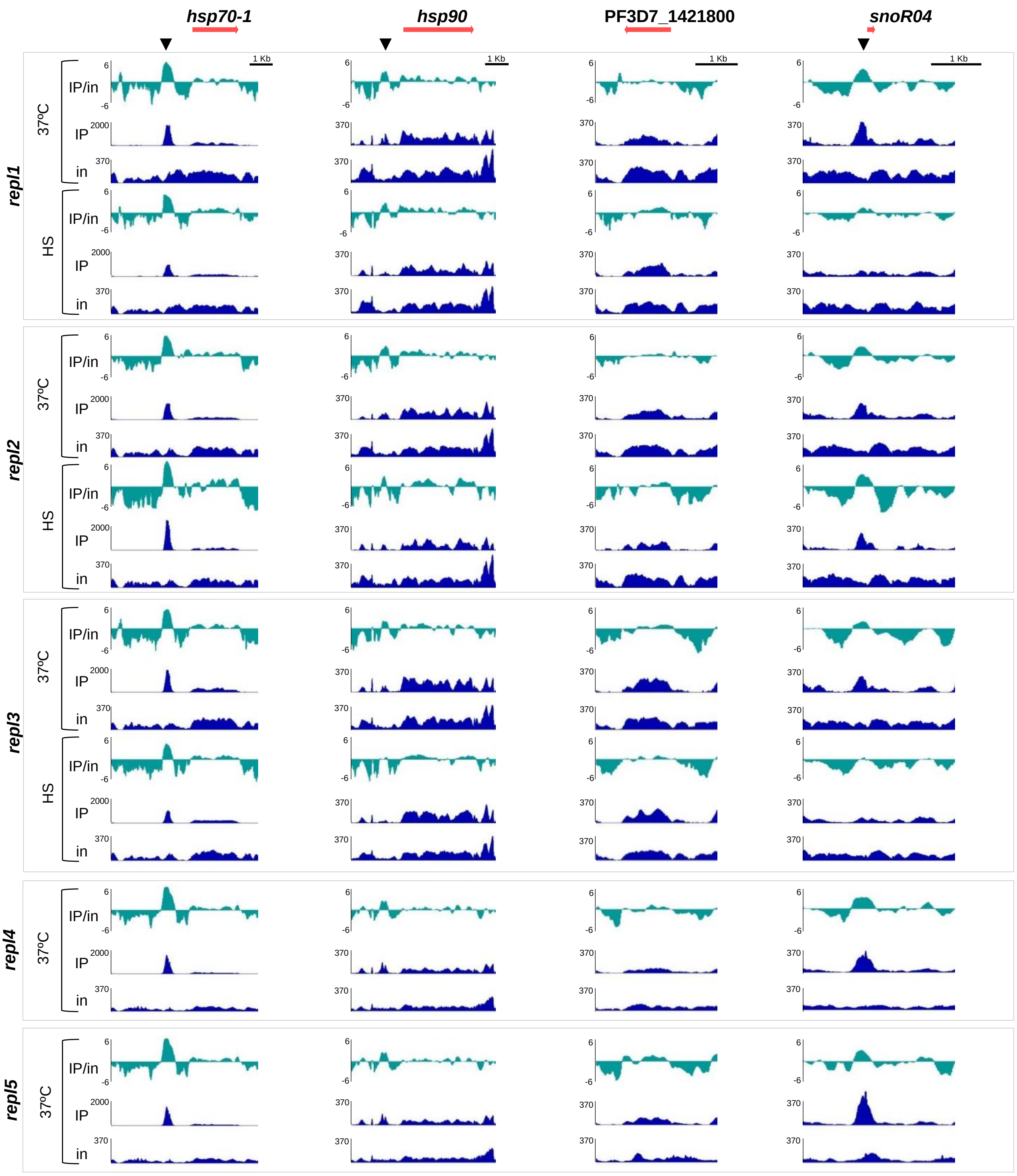


**b**

| | Chrom. | Start | End | Median MACS2 Score | Median Fold Enrich. | Closest Gene | Gene Name | Location |
|---|---|---|---|---|---|---|---|---|
| **Control** | **8** | 859109 | 859513 | 3155 | 14.6 | PF3D7_0818900 | *hsp70-1* | 5' |
| | **5** | 465548 | 465850 | 429 | 4.2 | PF3D7_0510900 | *snoR04* | 5' & CDS |
| | **13** | 2895119 | 2897998 | 150 | 2.3 | Telomere | - | -- |
| **HS** | **8** | 859123 | 859514 | 3410 | 21.8 | PF3D7_0818900 | *hsp70-1* | 5' |
| | **10** | 1437164 | 1437430 | 226 | 3.5 | PF3D7_1036400 | *lsa1* | CDS |
| | **13** | 2909970 | 2912982 | 181 | 2.9 | Telomere | - | - |

**c**

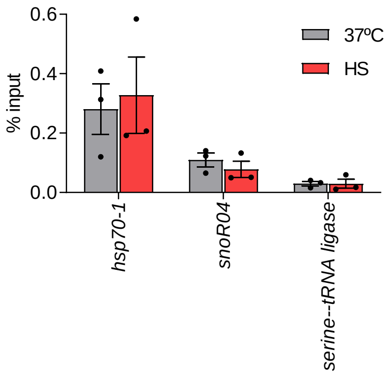

**Supplementary Figure 7. ChIP analysis of the chromosomal distribution of PfAP2-HS. a**, ChIP-seq analysis of HA-tagged PfAP2-HS. Number of reads of ChIP (IP) and input (in) tracks, and $log_2$-transformed ChIP/input ratio tracks (IP/in) for five independent biological replicates (three including HS and 37ºC conditions, two including only the 37ºC condition). Snapshots are shown for the three genes in cluster I (Fig. 2a) and *snoR04*. Binding at the *hsp70-1* and *hsp90* promoters coincides with the position of a tandem G-box motif, whereas PF3D7_1421800 and *snoR04* lack a G-box. **b**, Peaks present in ≥3 out of 5 replicate ChIP-seq experiments (37ºC) or ≥2 out of 3 replicate experiments (HS) and with a MACS score >100 in each positive replicate. **c**, ChIP-qPCR analysis of HA-tagged PfAP2-HS binding at selected loci, in cultures exposed to HS or control conditions (mean and s.e.m. of % input in *n*=3).

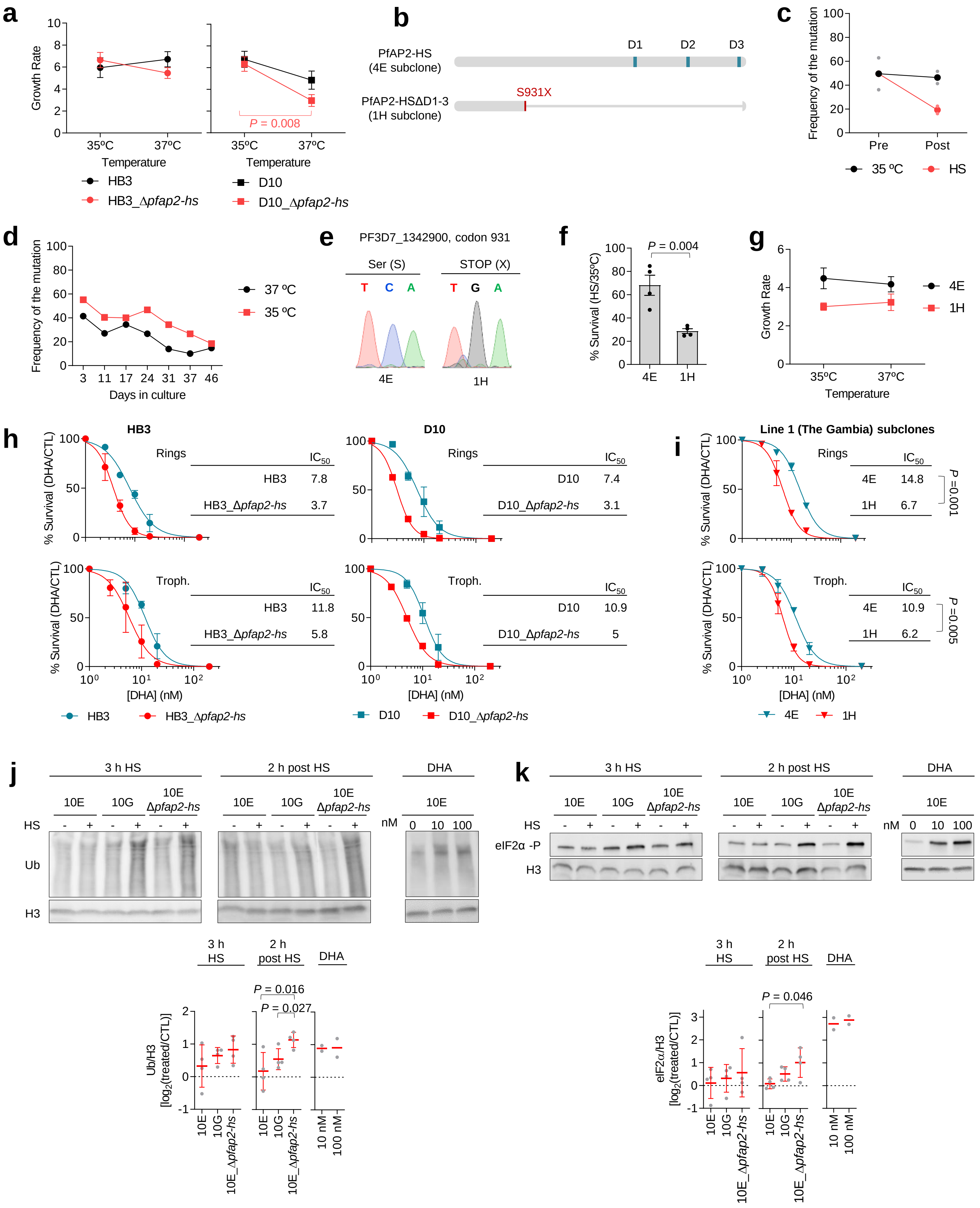
a
Growth Rate
35ºC 37ºC
Temperature
HB3
HB3_Δpfap2-hs
P = 0.008
D10
D10_Δpfap2-hs
b
PfAP2-HS (4E subclone)
D1 D2 D3
PfAP2-HSΔD1-3 (1H subclone)
S931X
c
Frequency of the mutation
Pre Post
35 ºC
HS
d
Frequency of the mutation
Days in culture
37 ºC
35 ºC
e
PF3D7_1342900, codon 931
Ser (S)
STOP (X)
T C A
T G A
4E
1H
f
% Survival (HS/35ºC)
P = 0.004
4E 1H
g
Growth Rate
35ºC 37ºC
Temperature
4E
1H
h
HB3
Rings
IC50
HB3 7.8
HB3_Δpfap2-hs 3.7
D10
Rings
IC50
D10 7.4
D10_Δpfap2-hs 3.1
% Survival (DHA/CTL)
Troph.
IC50
HB3 11.8
HB3_Δpfap2-hs 5.8
Troph.
IC50
D10 10.9
D10_Δpfap2-hs 5
[DHA] (nM)
HB3
HB3_Δpfap2-hs
D10
D10_Δpfap2-hs
i
Line 1 (The Gambia) subclones
Rings
IC50
4E 14.8
1H 6.7
P =0.001
Troph.
IC50
4E 10.9
1H 6.2
P =0.005
4E
1H
j
3 h HS
2 h post HS
DHA
10E 10G 10E Δpfap2-hs
HS - + - + - +
nM 0 10 100
Ub
H3
3 h HS
2 h post HS
DHA
P = 0.016
P = 0.027
Ub/H3 [log2(treated/CTL)]
10E 10G 10E_Δpfap2-hs
10 nM 100 nM
k
3 h HS
2 h post HS
DHA
10E 10G 10E Δpfap2-hs
HS - + - + - +
nM 0 10 100
eIF2α -P
H3
P = 0.046
eIF2α/H3 [log2(treated/CTL)]
10E 10G 10E_Δpfap2-hs
10 nM 100 nM

**Supplementary Figure 8. Phenotypic and transcriptional characterization of parasites lacking PfAP2-HS in different genetic backgrounds. a**, Growth rate of Δ*pfap2-hs* and parental lines at different temperatures (mean and s.e.m. of *n*=4). **b**, Schematic of wild-type PfAP2-HS and PfAP2-HS_ΔD1-3 observed in Line 1 from The Gambia as a consequence of a C to G mutation at codon 931 (S931X) arising during culture adaptation. The position of the AP2 domains is indicated (D1-3). **c**, Frequency of the mutation (as determined by Sanger sequencing) in culture-adapted Line 1 before (Pre) and after (Post) performing a HS at the trophozoite stage and culturing for an additional cycle (mean of *n*=2). **d**, Frequency of the mutation during additional time in culture at different temperatures. Day 0 is the day in which the frozen stock from The Gambia (culture-adapted for 91 days) was placed back in culture. **e**, Sanger sequencing confirmation of the presence or absence of the mutation at codon 931 in Line 1 subclones 4E and 1H. **f**, HS survival of 4E and 1H cultures. Control cultures were maintained at 35ºC (mean and s.e.m. of *n*=4). **g**, Growth rate of 4E and 1H at different temperatures (mean and s.e.m. of *n*=5). **h-i**, Survival (%) after a 3 h dihidroartemisinin (DHA) pulse at the ring or trophozoite stage. Values are the mean and s.e.m. of *n*=2 (**h**) or *n*=3 (**i**). **j-k**, Western blot analysis (representative of *n*=4) of polyubiquitinated proteins (Ub) (**j**) or phosphorylated eIF2α (eIF2α-P) (**k**) immediately after HS (3 h HS) and 2 h later (2 h post HS). Histone H3 is a loading control. DHA was used as a positive control, as it is a known inducer of the UPR[5,6]. The $Log_2$ of H3-normalized signal in HS or DHA-treated cultures versus control cultures is shown (mean and s.e.m. of *n*=4, for the DHA control mean of *n*=2). For gel source data, see Supplementary Fig. 11.

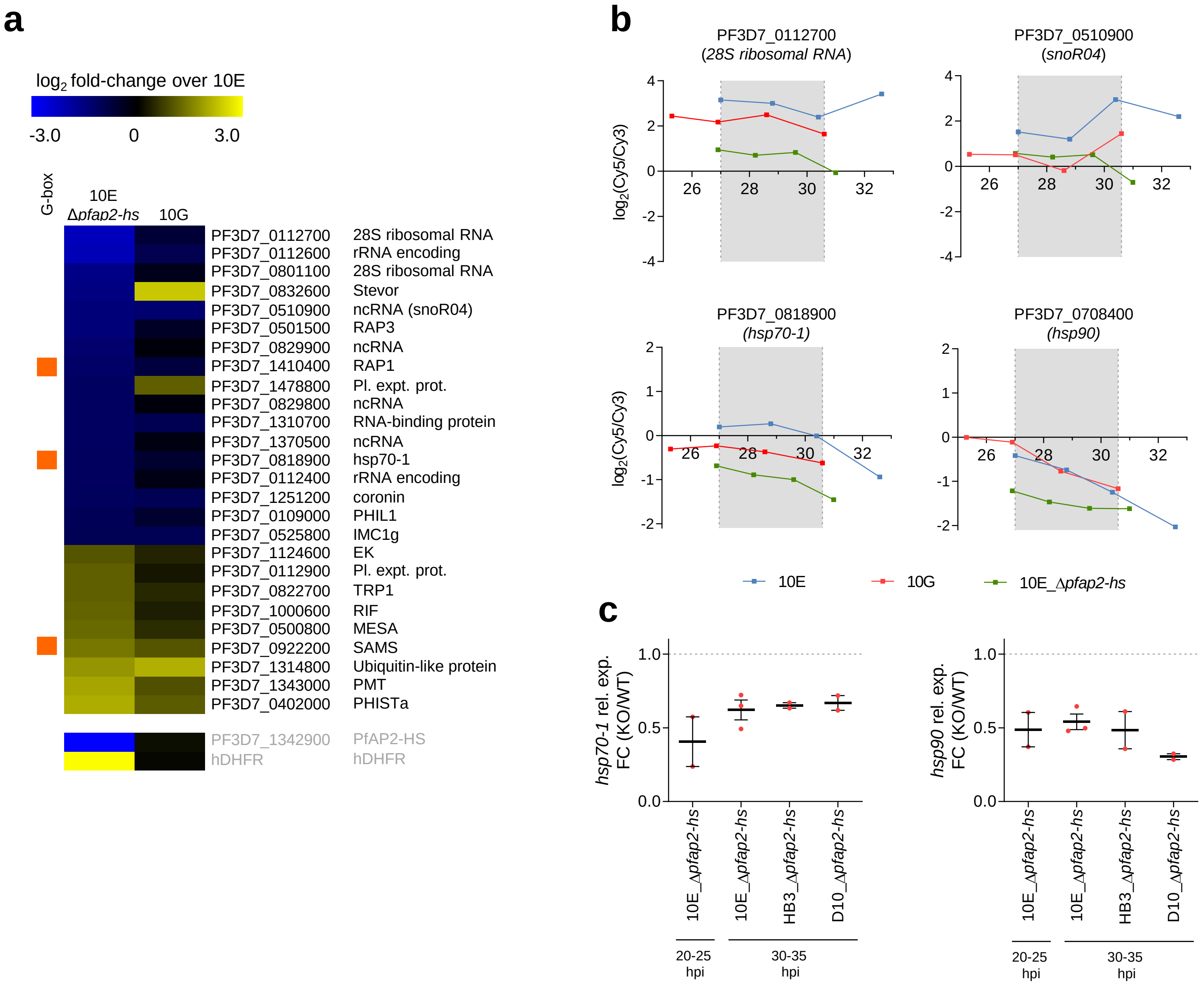


**Supplementary Figure 9. Transcriptional changes associated with PfAP2-HS deletion under basal (no heat shock) conditions. a**, Changes in transcript levels in the absence of HS for genes with an average expression fold-change >2 between 10E_Δ*pfap2-hs* and 10E. Values are the $\log_2$ of the average expression fold-change relative to 10E across the time period compared (~27-30.5 hpi). Genes artificially modified in the knockout line, which serve as controls, are shown at the bottom (their values are out of the range displayed). The column at the left indicates the presence of the G-box[3]. **b**, Expression plots for selected genes under basal conditions. Expression values are plotted against statistically-estimated parasite age, expressed in h post-invasion (hpi). Grey shading marks the interval used to calculate the average expression fold-change. **c**, RT-qPCR analysis of *hsp70-1* and *hsp90* transcript levels in *pfap2-hs* knockout lines compared to their wild type controls under basal conditions. Expression values are normalized against *serine--tRNA ligase,* and expressed as the fold-change (FC) in the knockout versus control lines. The mean and s.e.m. of *n*=3 (10E, 30-35 hpi) or *n*=2 (others) independent biological replicates is shown.

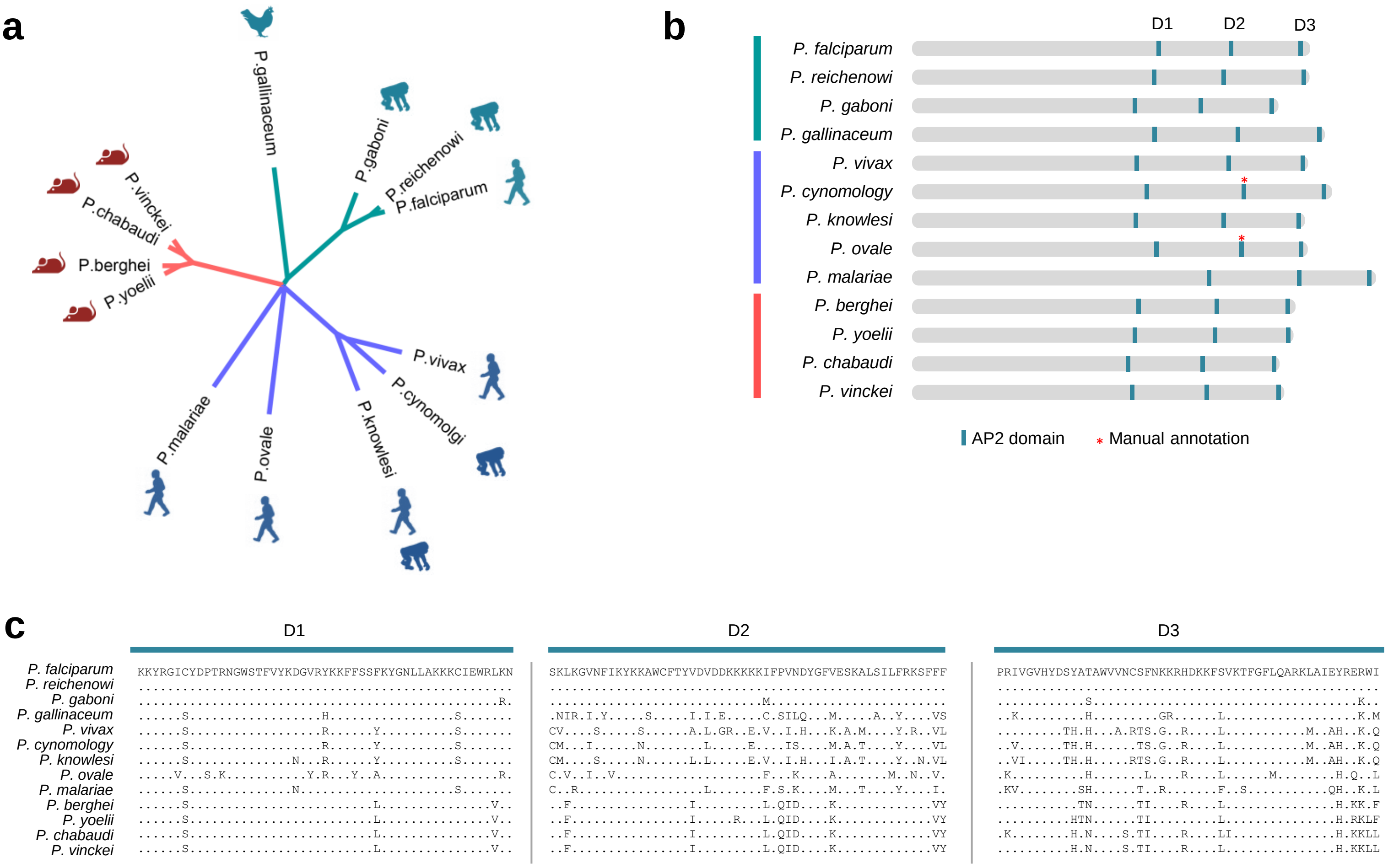


**Supplementary Figure 10. Phylogenetic analysis of AP2-HS. a**, Phylogenetic analysis of the protein sequence of AP2-HS orthologs in *Plasmodium* spp. The analysis includes one strain of each species for which the sequence is available in PlasmoDB. The sequences alignment and phylogenetic tree were constructed using Clustal Omega[7], with default parameters. A Neighbor-Joining tree without distance corrections was obtained. The cladogram was generated using FigTree 1.4.4. **b**, Schematic of the domain structure of AP2-HS orthologs in *Plasmodium* spp. The position of the AP2 domains is based on domains identified in PlasmoDB, except for those marked with an asterisk that were annotated manually according to the sequence alignment. **c**, Sequence alignment of the three AP2 domains (D1-D3) present in AP2-HS orthologs in *Plasmodium* spp. Dots indicate identity with the amino acid in the first sequence

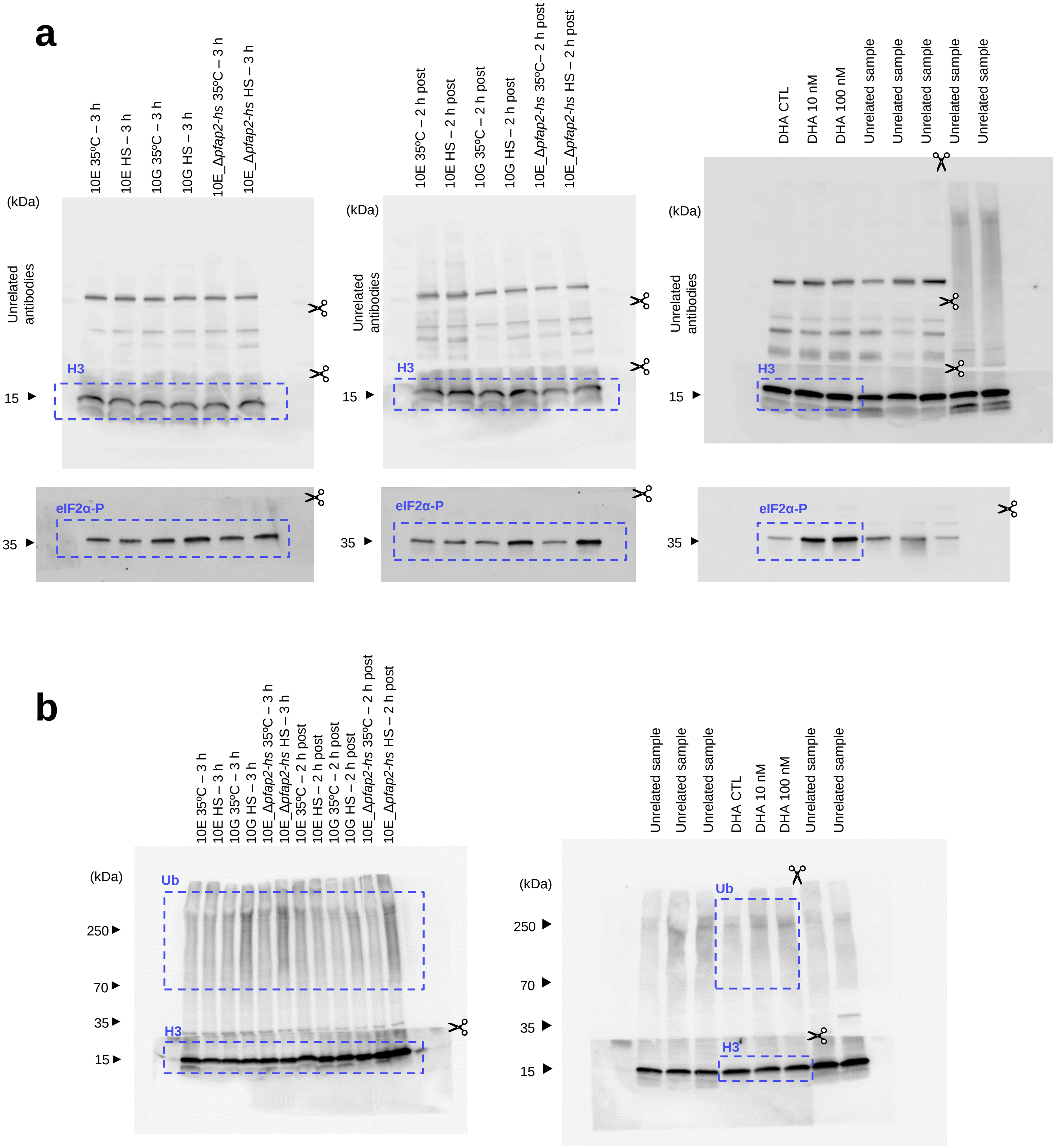


**Supplementary Figure 11. Gel source images for Western Blots.** Dashed blue lines indicate the position of the cropped images shown in Supplementary Fig. 8j-k. For quantification, lower exposure images were used. **a,** Membranes for eIF2α-P analysis were cut into 3 pieces (<25 kDa, ~25-55 kDa and >55 kDa) and tested for histone H3 (H3) loading control (<25 kDa piece) and other unrelated antibodies. Membranes for the ~25-55 kDa molecular weight range were stripped and reproved for eIF2α-P analysis. Therefore, the loading control was run in the same gel as eIF2α-P. **b,** Membranes for the analysis of polyubiquitinated proteins (Ub) were cut into 2 pieces (<25 kDa and >25 kDa) and tested for H3 and Ub, respectively. Therefore, the loading control was run in the same gel as Ub.
